# mDia2 is an important mediator of MRTF-A-dependent regulation of breast cancer cell migration

**DOI:** 10.1101/2023.12.21.572883

**Authors:** Ian Eder, Virginia Yu, Jacob Antonello, Fangyuan Chen, David Gau, Pooja Chawla, Marion Joy, Peter Lucas, David Boone, Adrian V. Lee, Partha Roy

**Affiliations:** Bioengineering, University of Pittsburgh; School of Medicine, University of Pittsburgh; School of Medicine, Tsinghua University, China; Hillman Cancer Center, University of Pittsburgh; Pathology, University of Pittsburgh; Biomedical Informatics, University of Pittsburgh; Pharmacology, University of Pittsburgh

**Keywords:** MRTF, actin, SRF, breast cancer, cell motility, diaphanous

## Abstract

Dysregulated actin cytoskeleton gives rise to aberrant cell motility and metastatic spread of tumor cells. This study evaluates the effect of overexpression of wild-type vs functional mutants of MRTF-A on migration and invasion of breast cancer (BC) cells. Our studies indicate that SRF’s interaction is critical for MRTF-A-induced promotion of both 2D and 3D cell migration, while the SAP-domain function is important selectively for 3D cell migration. Increased MRTF-A activity is associated with more effective membrane protrusion, a phenotype that is attributed predominantly to SRF’s interaction of MRTF. We demonstrate formin-family protein mDia2 as an important mediator of MRTF-stimulated actin polymerization at the leading edge and cell migration. Multiplexed quantitative immunohistochemistry and transcriptome analyses of clinical BC specimens further demonstrate a positive correlation between nuclear localization of MRTF with malignant traits of cancer cells and enrichment of MRTF-SRF gene signature in pair-matched distant metastases vs primary tumors. In conclusion, this study establishes a novel mechanism of MRTF-dependent regulation of cell migration and provides evidence for the association between MRTF activity and increased malignancy in human breast cancer, justifying future development of a specific small molecule inhibitor of the MRTF-SRF transcriptional complex as a potential therapeutic agent in breast cancer.

**SIGNIFICANCE:**

- Actin cytoskeletal dysregulation gives rise to metastatic dissemination of cancer cells. This study mechanistically investigates the impact of specific functional disruption of MRTF (a transcriptional co-factor of SRF) on breast cancer cell migration.
- This study establishes a novel mechanism linking mDia2 to MRTF-dependent regulation of cell migration and provides clinical evidence for the association between MRTF activity and increased malignancy in human breast cancer.
- Findings from these studies justify future exploration of specific small molecule inhibitor of the MRTF-SRF transcriptional complex as a potential therapeutic agent in breast cancer.

## INTRODUCTION

Most breast cancer (BC)-related deaths are due to the metastatic spread of tumor cells to distant organs. Metastasis is a complex process consisting of multiple steps, including tumor cell invasion through the stromal tissue, intravasation, extravasation, and ultimately colonization of tumor cells at the distant organ site. The initial step in this cascade of events, stromal invasion of cancer cells, is highly dependent upon dynamic actin remodeling in tumor cells and the action of various secreted ECM-degrading proteases. Disruption of actin cytoskeleton homeostasis is a critical feature of metastatic cancer cells. In principle, restoration of actin cytoskeleton normality should attenuate the migratory propensity of tumor cells thereby reducing metastatic burden in BC patients.

Myocardin-related Transcription Factors (MRTFs), a family of transcriptional co-activators, are major regulators of the expression of genes relevant to cytoskeleton organization, cell adhesion and ECM remodeling (summarized in (Gau and Roy 2018)). There are two isoforms of MRTF in non-muscle cells: MRTF-A (also known as MKL1 or MAL), and MRTF-B (also known as MKL2). Both isoforms of MRTF share common structural motifs. At the N-terminus, they contain three RPEL domains which bind to G-actin as a pentavalent G-actin-MRTFA complex (Mouilleron, et al. 2011). In between the various RPEL domains, there are spacer regions, and interspaced between the last two RPEL domains, there is a bi-partite basic-rich region (B2-B3) containing a nuclear localization signal (NLS) which becomes exposed once G-actin is unbound (Nakamura, et al. 2010; Pawlowski, et al. 2010; Mouilleron, et al. 2011). The other conserved domains are a basic-rich B1 domain (contains a second NLS and SRF-binding region), a glutamine-rich Q domain (involved in SRF’s interaction and nuclear export), a SAP domain (a putative chromatin-binding domain), a leucine-zipper domain (mediates homo- and hetero-dimerization), a PPxY domain (mediates interactions with WW domains of partner proteins including YAP (Yes-associated protein)), and a C-terminal transcriptional activation domain (Miralles, et al. 2003; Mouilleron, et al. 2011).

MRTF is a highly regulated transcription factor. In resting cells, binding to G-actin sequesters MRTF in the cytoplasmic compartment restricting its transcriptional activity. Induction of actin polymerization upon growth-factor stimulation liberates MRTF from G-actin, allowing MRTF to translocate to the nucleus and participate in gene regulatory activities. Therefore, nuclear translocation and transcriptional activity of MRTF are highly sensitive to the relative abundance of G- and F-actin in cells (Miralles, et al. 2003; Posern, et al. 2004; Vartiainen, et al. 2007). When in the nucleus, MRTF directly binds to SRF orchestrating a broad transcriptional program including expression of SRF itself and many other genes, a subset of which are relevant for actin cytoskeletal control (Gualdrini, et al. 2016). Notably, MRTF preferentially interacts with SRF as a dimer rather than a monomer (Miralles, et al. 2003). Although MRTF is best known for its role as a transcriptional co-factor of SRF, it is hypothesized that MRTF is also capable of gene regulation in an SRF-independent manner directly utilizing its SAP domain (Gurbuz, et al. 2014; Asparuhova, et al. 2015).

A number of previous studies have demonstrated that loss of expression of either MRTF isoforms or SRF causes defects in actin cytoskeleton and cell migration in physiological and pathological contexts (summarized in (Gau and Roy 2018)). However, since MRTF has SRF-independent gene regulatory activities, and SRF has MRTF-independent gene regulatory activities through its binding with ternary complex factors (TCFs) (Hill, et al. 1993; Gualdrini, et al. 2016), the precise role of MRTF-SRF interaction in cell migration cannot be extrapolated from those loss-of-function studies. In this study, we for the first time directly demonstrate the critical importance of MRTF-SRF interaction in cell migration and invasion, link upregulation of formin-family actin-binding protein (ABP) mammalian Diaphanous-2 (mDia2 – encoded by the *DIAPH3* [diaphanous-related formin 3] gene) expression as a mechanism of MRTF-directed control of cell migration, and further provide clinical evidence for MRTF activity association with increased malignancy in human BC.

## RESULTS

### MRTF promotes migration and invasion of breast cancer cells requiring its SRF’s interaction

To investigate the role of the MRTF-SRF interaction in breast cancer cell motility, we performed cell migration studies using MDA-MB-231 (MDA-231), a widely used human metastatic triple-negative breast cancer (TNBC - lacks the expression of estrogen receptor (ER), progesterone receptor (PR) and HER2 receptor) cell line as a model system, in an overexpression (OX)-based experimental setting. A previous study found that MDA-231 cells exhibit reduced migration (in a wound-healing assay) and invasion *in vitro* and *in vivo* when MRTF isoforms are stably co-silenced (Medjkane, et al. 2009), demonstrating MRTF-dependence for motility of MDA-231 cells and justifying its use as a model cell line for the purpose of our studies.

In our overexpression studies, we utilized our previously generated GFP/luciferase (Luc) expressing sublines of MDA-231 cells engineered for doxycycline (dox)-inducible stable overexpression of either wild-type MRTF-A (WT-MRTF-A), a SAP-domain-deleted mutant form of MRTF-A (referred to as ΔSAP-MRTF-A herein), or an SRF-binding-deficient point-mutant of MRTF-A (referred to as ΔSRF-MRTF-A herein) (Gau, et al. 2022). A dox-treated GFP/Luc-expressing subline without any additional MRTF-A perturbation served as the control. **Fig 1A** depicts the mutations performed to generate the ΔSRF-MRTF-A and ΔSAP-MRTF-A constructs. The four point mutations (K237A, Y238A, H239A, Y241A) in the B1 domain of MRTF-A were designed based on the findings of alanine linker-scanner mutagenesis studies that assessed the complex formation between a truncated constitutively active MRTF-A, DNA-binding domain of SRF, and a synthetic SRF-binding DNA probe in gel-shift mobility assays (Zaromytidou, et al. 2006). We also performed a pull-down assay with HEK-293 cell lysates to confirm that alanine substitutions on three of these four residues (K237/Y238/H239) are sufficient to dramatically abrogate MRTF-A’s binding to recombinant GST-SRF (**Fig 1B**). As per our RNA-sequencing data (discussed later), the transcript level of MRTF-A was relatively comparable across the various MRTF-A overexpressing cell lines. However, at the protein level, the total MRTF-A expression was demonstrably lower in ΔSRF-MRTF-A overexpressing cells relative to either WT- or ΔSAP-MRTF-A overexpressing sublines even though the expression level of mCherry (a reporter co-expressed with the various MRTF-A constructs under the same dox-inducible promoter in an IRES system) was similar across the various sublines (**Fig 1C** and **Fig S1).** Since the antibody we used to detect MRTF-A expression targets an epitope (residues surrounding G812) in the C-terminal TAD domain that is far away from either the B1 or the SAP-domain of MRTF-A, we do not think that the lower MRTF-A expression of ΔSRF-MRTF-A is an artifact of an antibody recognition issue. Based on comparable transcript levels of MRTF-A across the various overexpression sublines, it is also unlikely that MRTF-A expression is transcriptionally augmented in a positive feedback manner when either WT- or ΔSAP-MRTF-A is exogenously overexpressed and blocked when SRF’s interaction of MRTF-A is disrupted. However, it is possible that disrupting SRF’s interaction of MRTF-A may somehow impact the protein translation and/or the rate of protein turnover of MRTF-A. Pertaining to the latter, a previous study reported that FHL2 (four and half Lim-domain protein 2) interacts with both isoforms of MRTF and that FHL2 promotes the expression of MRTF-A at the protein but not at the mRNA level mainly through increasing its protein stability (Hinson, et al. 2008). Although the exact residues of MRTF-A:FHL2 interaction have not been mapped, regions containing the B1 and B2 domains of MRTF-A are required for this interaction. Given these findings, it is highly plausible that ΔSRF-MRTF-A has a higher rate of protein turnover than WT-MRTF-A potentially because of a reduced affinity for FHL2.

**Figure 1.**
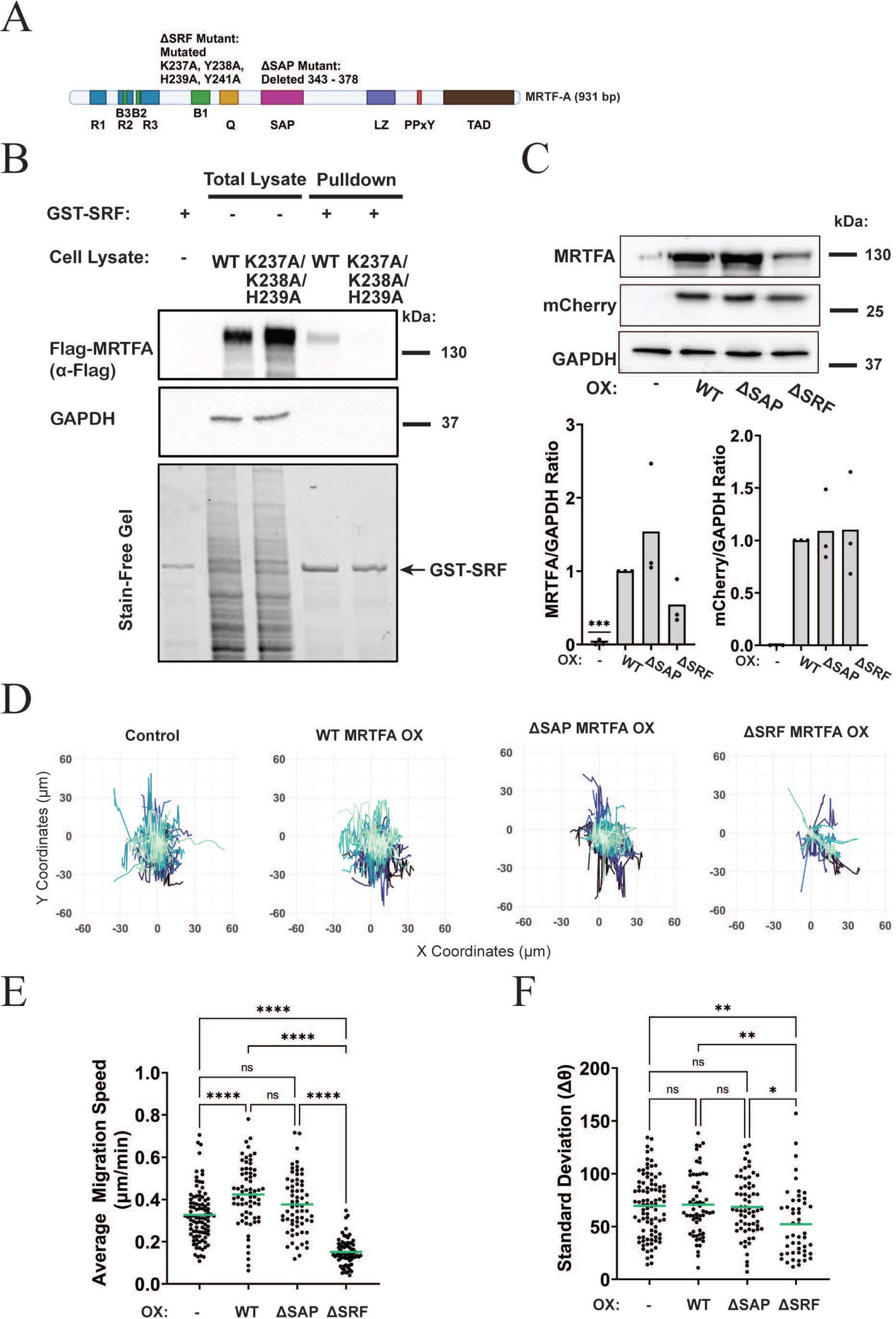
Effect of overexpression of WT vs mutant forms of MRTF-A on 2D random motility of breast cancer cells. A) Schematic of the various functional domains and the mutations created to generate the ΔSAP- and ΔSRF-mutants of MRTF-A. **B)** Representative immunoblots and associated quantification (n=3 experiments) demonstrating dox-induced overexpression (OX) of wild-type (WT) and the various mutant forms of MRTF-A along with mCherry (reporter) in GFP/luciferase expressing subline of MDA-231 cells (GAPDH serves as a loading control). One-sample T-test was used to determine significance. *** indicates p value < 0.001. **C)** Left: Anti-flag immunoblot of total lysates and GST-SRF pulldown samples of HEK-293 lysate transiently transfected with flag-tagged WT- and K237A/K238A/H239A-flag-tagged MRTF-A plasmids (GAPDH blot serves as the loading control of total cell lysates); Right – Stain-free gel images show comparable levels of GST-SRF for the two pull-down conditions. **D-E)** Trajectories of individual MDA-231 cells of different experimental groups in 2D time-lapse motility assay performed after 48 hours of dox treatment (*panel D*). Dot plots (with indicated means) in *panels E* and *F* represent the average migration speed and the standard deviation of Δθ (a measure of the change in direction), respectively, pooled from three independent experiments (control: n = 96 cells, WT-MRTF-A OX: n = 66 cells, ΔSAP-MRTF-A OX: n = 65 cells, and ΔSRF-MRTF-A OX: n = 69 cells). Outliers were removed using the ROUT method (Q = 1%). One-way ANOVA was performed. **** indicates p value < 0.0001, ** indicates p value < 0.01, and * indicates p value < 0.05.

Although we previously showed that deletion of the entire B1 domain reduces the nuclear localization of MRTF-A (this is consistent with the presence of an NLS in the B1 domain - (Joy, et al. 2017)), we confirmed that select point-mutations in the B1 domain to disrupt SRF’s interaction does not negatively impact the nuclear localization of MRTF-A. In fact, our immunostaining data unexpectedly revealed a much more striking nuclear presence of MRTF-A in ΔSRF-MRTF-A expressers relative to the other three sublines (**Fig S2**). Even though these immunostaining data reflect both endogenous and exogenous MRTF-A isoforms, the expression of endogenous MRTF-A is dramatically lower than any of the exogenous forms of MRTF-A (including ΔSRF-MRTF-A). Therefore, it is reasonable to presume that the strong nuclear localization of MRTF-A in ΔSRF-MRTF-A expressers is largely reflective of the mutant form of MRTF-A. While the underlying reasons are not completely clear, at least, this assured us that the phenotypic changes induced by ΔSRF-MRTF-A overexpression would not be a simple consequence of failure in nuclear localization of the mutant MRTF-A.

To investigate the effect on migration of the function inhibited mutants of MRTF-A, we first performed time-lapse single-cell imaging of 2D random motility of the various dox-inducible MRTF-A overexpressing MDA-231 sublines with transgene overexpression turned on 48h prior to time-lapse imaging. **Fig 1D** shows the trajectories of different groups of cells obtained from 3-hour time-lapse recordings (pooled from three independent experiments). Consistent with slower 2D random motility of MDA-231 cells following acute co-silencing of MRTF isoform (**Fig S3**), overexpression of WT-MRTF-A accelerated cell migration by ∼30% relative to control cells in a statistically significant manner (**Fig 1E**). Although overexpression of ΔSAP-MRTF-A resulted in a marginal 15% increase in the average migration speed relative to control cells, this difference was not statistically significant. The most striking feature from these experiments is a robust 54% decrease in the speed of migration exhibited by the ΔSRF-MRTF-A overexpressing cells when compared to control cells. We also analyzed the effect of overexpression of these MRTF-A variants on the persistence of cell migration. To represent this metric, we computed the standard deviation of the change in migration direction (Δθ) for each time step. Higher standard deviation of Δθ is interpreted as greater fluctuation in the direction of cell migration. While the values of standard deviation of Δθ were not statistically different between control, WT-MRTF-A expressers, and ΔSAP-MRTF-A expressers, overexpression of ΔSRF-MRTF-A improved the persistence of cell migration, as indicated by a lower standard deviation of Δθ compared to other three sublines in a statistically significant manner (**Fig 1F**). These findings demonstrate that SRF’s interaction is critical for MRTF’s ability to promote 2D cell migration.

To better represent the *in vivo* scenario of stromal invasion of cancer cells, we performed two types of invasion experiments with these cell lines. First, we analyzed the random single-cell invasion speed within a sandwiched 3D collagen-I matrix with transgene overexpression turned on 48h prior to time-lapse imaging. **Fig 2A** shows the trajectories of different groups of cells obtained from a 12-hour time lapse recording (pooled from three independent experiments). Consistent with the trends of 2D random migration experiments, overexpression of WT-MRTF-A increased the average 3D cell migration speed by ∼16%, while overexpression of ΔSRF-MRTF-A decreased cell migration speed by ∼36% as compared to the control group of cells (**Fig 2B**). Although the average migration speed of ΔSAP-MRTF-A expressers was not statistically different from that of control cells in 2D random migration assay, ΔSAP-MRTF-A expressers exhibited slower (by ∼25%) random invasion speed relative to control in a statistically significant manner in a 3D ECM environment (**Fig 2B**). We also analyzed the directionality of movement (measured as a cell’s end-to-end distance divided by the total distance traveled by the cell) across different groups. As per these measurements, the directionality readouts of all three MRTF-A overexpression groups were lower than that measured for the control group (**Fig 2C**). Next, to specifically mimic directed invasion of cancer cells from a tumor, we performed tumor spheroid invasion assays in collagen-I matrix over a period of 96 hours. Since we previously found that induction of ΔSRF-MRTF-A expression inhibits tumor spheroid formation, dox-induced transgene expression for all experimental groups was turned on only after the spheroid formation was complete. Representative images of tumor spheroids for different groups at the 0- and 96-hour time-points (by either phase-contrast imaging or fluorescence imaging of DAPI/phalloidin-stained cells) are shown in **Fig 2D**. To account for differences in tumor spheroid growth, we normalized the endpoint DAPI positive area to the initial spheroid area (assessed from the phase-contrast images) as a metric for directed invasion of tumor spheroids. Consistent with our 3D and 2D single-cell migration data, directed collagen invasiveness of ΔSRF-MRTF-A expressing tumor spheroids was also strikingly lower (by 69%) than the control spheroids (**Fig 2E**). Although in a random 3D migration assay, overexpression of WT-MRTF-A slightly enhanced the absolute speed of invasion of individual cells relative to control, the directed invasion readout of WT-MRTF-A-expressing tumor spheroids was lower than the control group. We speculate that this paradoxical observation is most likely due to the negative impact of MRTF-A overexpression on the directional persistence of movement (as demonstrated in **Fig 2C**). Consistent with single-cell invasion data, overexpression of ΔSAP-MRTF-A also dramatically reduced the directed invasion of tumor spheroids (**Fig 2E**). There are two key take-home messages from these 2D and 3D migration experiments. First, MRTF-A promotes the intrinsic 2D and 3D cell migration requiring its interaction with SRF. However, the dependence on the SAP-domain related activity of MRTF-A is likely more pertinent for cell migration in a 3D environment. Second, an excessively high level of MRTF-A activity may be somewhat counterproductive for directed tumor cell invasion. Why in some cases overexpression of MRTF-A mutants not only reversed the hypermigration phenotype of WT-MRTF-A overexpressing cells but also actively inhibited cell migration is discussed later.

**Figure 2.**
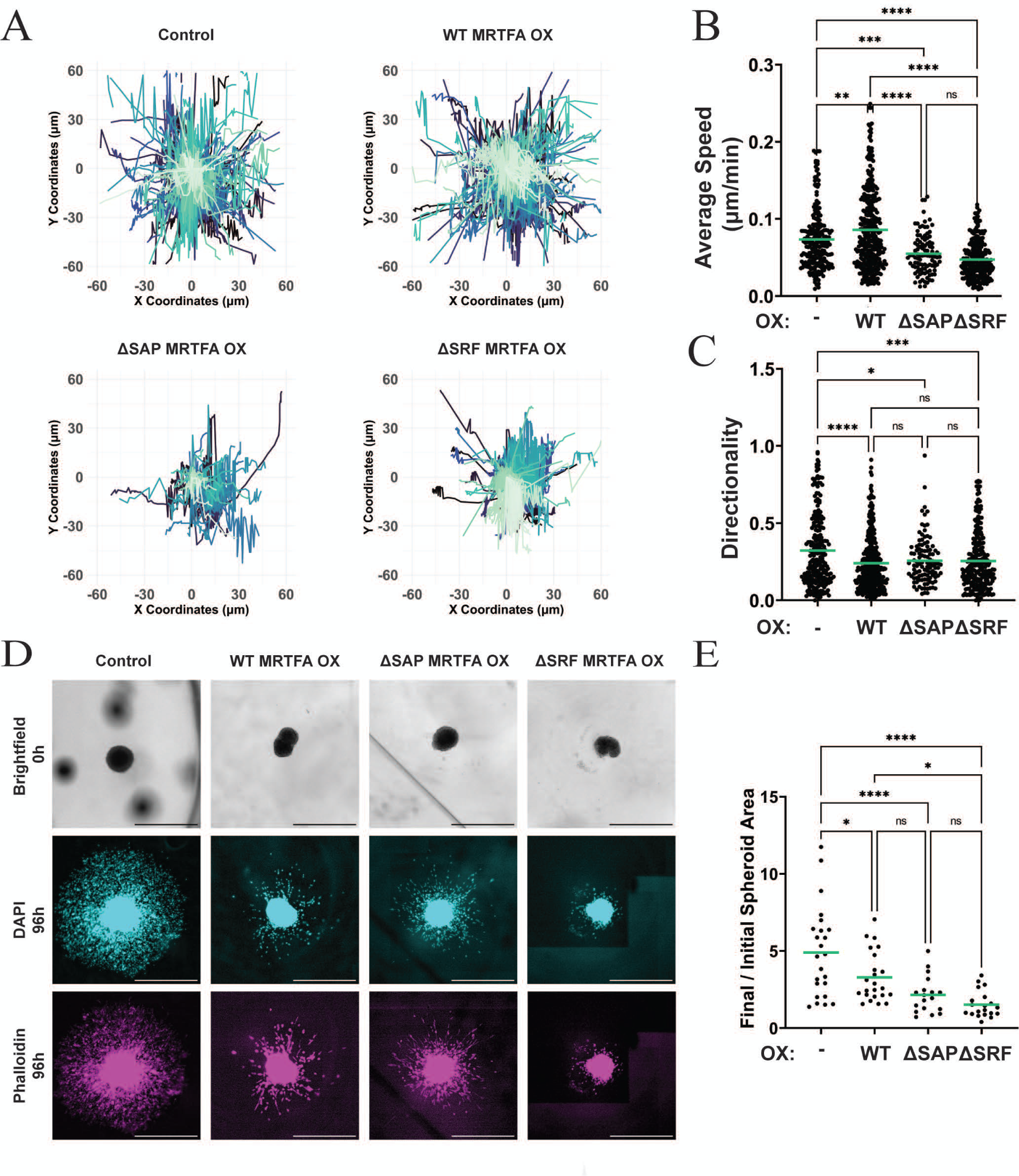
Effect of overexpression of WT vs mutant forms of MRTF-A on 3D single cell invasion and directed invasion of tumor spheroids A) Trajectories of individual MDA-231 cells of different experimental groups in a 3D time-lapse invasion assay. **B, C)** Dot plots (with indicated means) of the average invasion speed (*panel B*) and directionality (a measure of persistent of migration direction) (*panel C*). Data was pooled from four independent experiments (control: n = 223 cells, WT-MRTF-A OX: n = 348 cells, ΔSAP-MRTF-A OX: n = 100 cells, ΔSRF-MRTF-A OX: n = 240 cells; OX -overexpression). **D)** Representative images of MDA-231 tumor spheroids embedded in 1mg/ml type 1 collagen matrix at 0h (phase-contrast – 2.5X) and 96h (fluorescence staining: DAPI – 2.5X, Phalloidin – 2.5X). **E)** Quantification of the final invasive area as measured by the total DAPI+ region after 96h normalized to the size of the spheroids at 0h as measured from the phase-contrast images (control: n = 24 spheroids, WT-MRTFA OX: n = 23 spheroids, ΔSAP-MRTF-A OX: n = 18 spheroids, ΔSRF-MRTF-A OX: n = 20 spheroids). Spheroid invasion data were pooled from 7 independent experiments. Outliers were removed using the ROUT method (Q = 1%). One-way ANOVA was performed. **** indicates p value < 0.0001, ** indicates p value < 0.01, and * indicates p value < 0.05. Scale bar = 900µm.

### SRF’s interaction is important for MRTF-dependent regulation of membrane protrusion

Since lamellipodial protrusion is the initiating step for cell migration, to further understand the physical basis for the defect in cell migration induced by the disruption of MRTF-SRF binding, we comparatively analyzed the leading-edge dynamics between the various groups of cells by performing kymography analyses. From the representative kymography scans of the various sublines of MDA-231 cells (**Fig 3A**), it was qualitatively obvious that unlike the other three sublines, ΔSRF-MRTF-A expressers exhibited much less fluctuation of the membrane (marked by cycles of protrusions and retraction) suggesting that membrane dynamics are suppressed upon inhibition of the MRTF-SRF interaction. In order to quantitatively describe these changes, we used an automated image analysis pipeline to detect local minima and maxima in the protruding membrane at the leading edge. Based on these analyses, we quantified several parameters including the frequency of protrusion, the duration of protrusion ascent (i.e. the time taken for the protrusion to go from the local minimum to the local maximum) and the net protrusion distance (i.e. equal to the protrusion distance minus the retraction distance), the results of which are summarized in **Figs 3B-D** and **Fig S4**. Although overexpression of WT-MRTF-A reduced the frequency of initiating protrusions relative to control, the duration of protrusion ascent significantly increased (**Figs 3B-C**) and these protrusions underwent less retraction (possibly reflecting better protrusion-adhesion coupling – this is not unexpected given MRTF’s role in regulating cell-adhesion (Kishi, et al. 2016))(**Fig S4**). The combination of these features improved the net protrusion distance (a critical requirement for effective cell motility) in WT-MRTF-A overexpressers *vs* control. Disruption of the SAP-domain interaction of MRTF-A did not reverse the MRTF-A-induced increase in either the net protrusion distance or the duration of protrusion ascent. However, inhibition of SRF’s interaction with MRTF-A resulted in reduction in the frequency and distance of protrusion (lowest among the four groups) and completely reversed the MRTF-A-induced increase in the net protrusion distance (**Figs 3B-C; Fig S4**). Interestingly, although protrusions were initiated much less frequently in ΔSRF-MRTF-A expressers, the average duration of protrusion ascent of these cells (18.68 ± 8.76s) was longer than that calculated for control cells (15.08 ± 7.01s), an observation that is consistent with the trend of improved persistence of motion exhibited by these cells as illustrated in **Fig 1E**.

**Figure 3.**
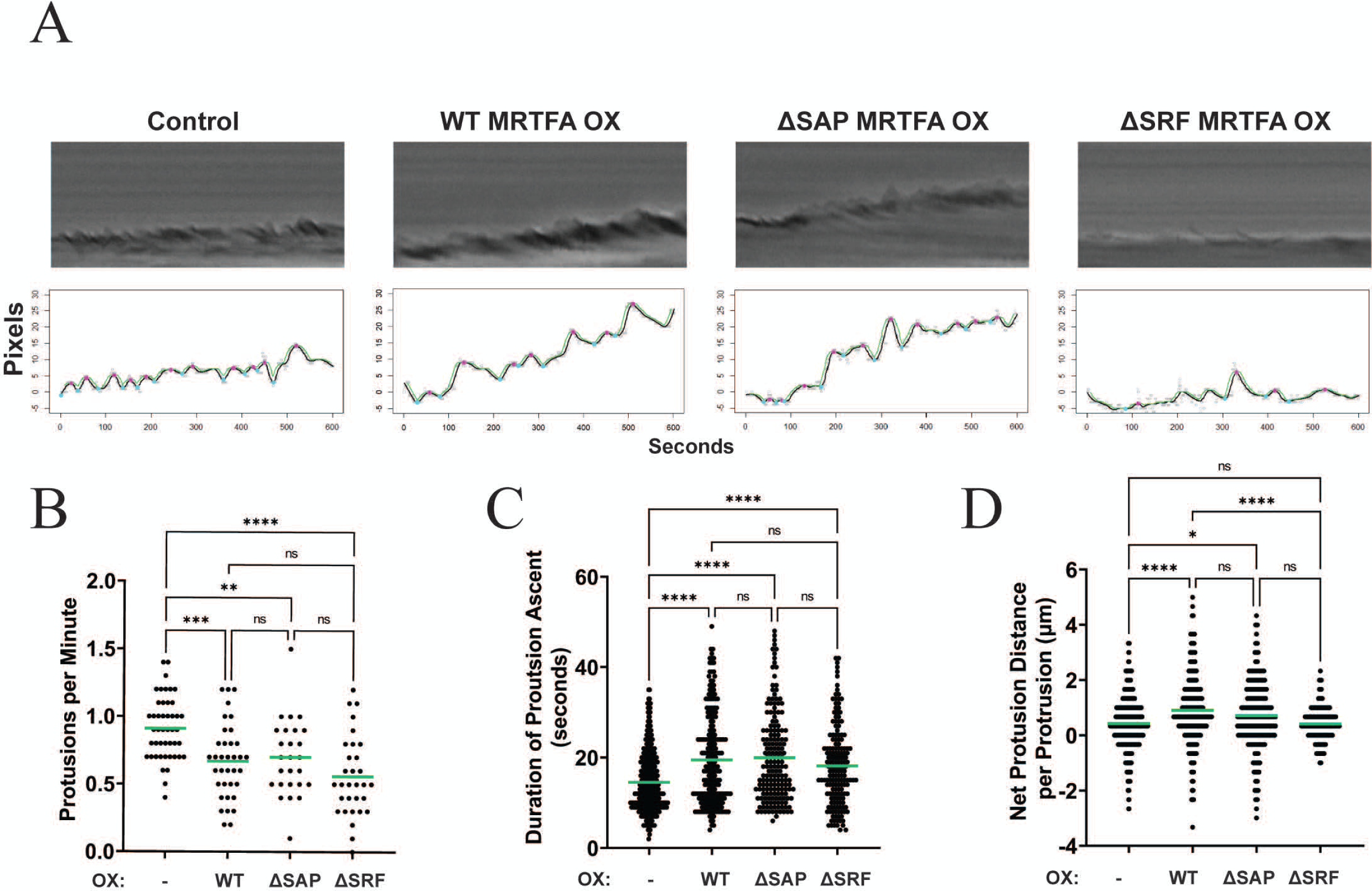
Effect of overexpression of WT vs mutant forms of MRTF-A on the features of lamellipodial protrusion. A) Representative kymographs constructed at the leading edge of migrating cells (time-lapse imaging was performed at 20X magnification for a total 10-minute duration at a 2-second interval) of the indicated groups of cells (OX – overexpression). The top and bottom rows are representative scans from phase-contrast images and the corresponding automated quantification of the peaks in the kymographs, respectively. The black traces represent a fitted smoothed curve calculated by local regression (LOESS) method. The green lines above the black lines represent the moving maximum point of the approximate leading front of the cells (as determined by the fitted smoothed curve) for the surrounding points (+/- 6 seconds). Local maxima indicated by the pink dots were counted when the black lines intersect the green lines. Reverse operations were performed to calculate the local minima indicated by the blue dots. **B)** Quantification of the number of protrusions per minute at each leading edge measured for the indicated groups (number of leading edges analyzed: control - 46, WT-MRTF-A OX - 36, ΔSAP-MRTF-A-OX - 26, and ΔSRF-MRTF-A OX - 31). **C)** Quantification of the net protrusion distance (extension distance – retraction distance) per protrusion for the indicated groups (number of protrusions analyzed: control -400, WT-MRTF-A OX –229, ΔSAP MRTFA OX -174, and ΔSRF MRTFA OX: n = 169 protrusions). **D)** Quantification of the duration of protrusion ascent per protrusion of the indicated groups (number of protrusions analyzed: Control - 395, WT-MRTF-A OX – 210, ΔSAP-MRTF-A OX -179, and ΔSRF-MRTF-A OX -159). Kymographs were pooled across multiple fields and wells in 3 independent experiments. For all quantifications, outliers were removed using the ROUT method (Q = 1%). One-way ANOVA was performed. **** indicates p value < 0.0001, *** indicates p value < 0.001, ** indicates p value< 0.01, and * indicates p value < 0.05.

We previously showed that MRTF-A overexpression induces a global increase in actin polymerization in MDA-231 cells requiring SRF’s interaction (Gau, et al. 2022). Given that actin polymerization at the leading edge is critical for lamellipodial protrusion, we next analyzed the phalloidin intensity at the lamellipodial edge of the various sublines of MDA-231 cells in a wound-healing assay to induce a polarized phenotype with well-defined lamellipodia in migratory cells at the edge of the wound (**Fig 4A**). Consistent with our previous findings related to MRTF-A’s effect on global F-actin content, overexpression of WT-MRTF-A also augmented actin polymerization at the leading edge (marked by an increase in the phalloidin intensity) that was completely reversed when SRF’s interaction but not the SAP-domain function of MRTF-A was disrupted (**Fig 4B**). Note that although ΔSRF-MRTF-A expressers exhibited slower migration speed than control cells, this difference did not translate when we compared the mean phalloidin intensity at the lamellipodial edge between these two groups of cells (**Fig 4B**). This prompted us to query whether disruption of SRF’s interaction impacts other types of F-actin-based protrusive structures. Since filopodia also play an important role in directional sensing and cell migration in both 2D and 3D environments (Jacquemet, et al. 2015), we quantitatively analyzed the relative filopodia formation abilities of these various sublines in response to acute stimulation of bradykinin (induces filopodia formation through activating Cdc42 family of Rho-GTPases (Kozma, et al. 1995)). Our experiments showed that overexpression of MRTF-A increased the average number of filopodia/cell as well as the average filopodial length relative to control (**Figs 4C-E**). Disrupting MRTF-A’s SRF interaction ability caused reversal of these changes, more prominently and statistically significantly for the average filopodial length. However, neither of these two parameters were significantly different between control and ΔSRF-MRTF-A expressers. Deletion of the SAP-domain had no negative impact on MRTF-A’s ability to augment filopodial abundance and only marginally reduced the average filopodial length relative to WT-MRTF-A overexpression group. Overall, these results indicate that MRTF-A-regulated actin-based membrane protrusion is dependent on its SRF’s interaction.

**Figure 4.**
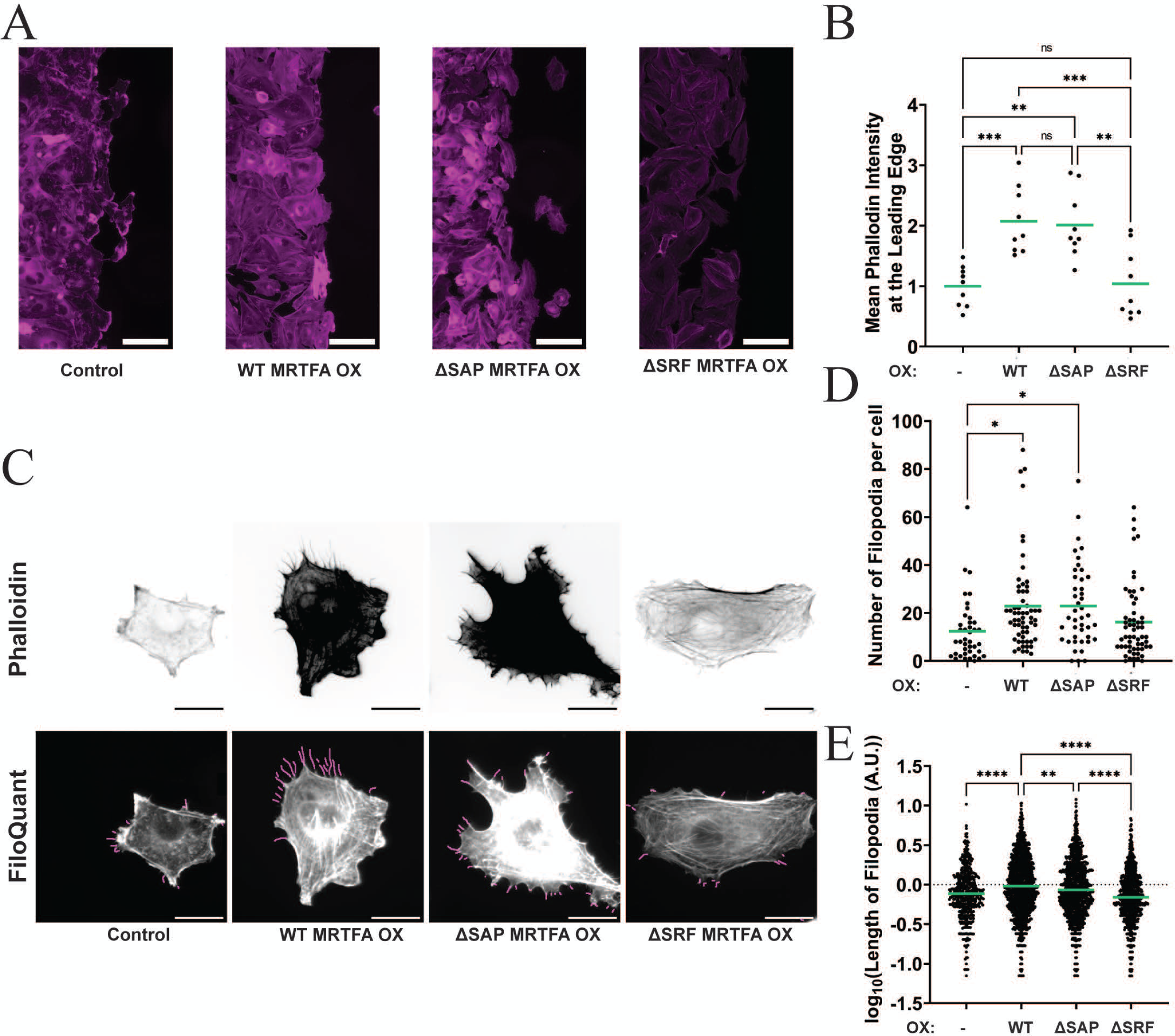
Effect of overexpression of WT vs mutant forms of MRTF-A on actin polymerization at the leading edge and filopodial formation. A, B) Representative phalloidin staining images (*panel A;* scale bar = 100 µm) at the migrating front of a wound-healing assay and quantification of mean phalloidin intensity at the leading edge of cells (*panel B*) (images were acquired 3 hours after the monolayer was scratched, and 9 leading fronts were analyzed/group. Data points are pooled from 3 independent experiments with 3 technical replicates per experiment. **C)** Representative phalloidin staining images of bradykinin stimulated MDA-231 cells with indicated perturbations of MRTF-A (top) and automated detection of filopodia by FiloQuant (bottom) (OX – overexpression). **D)** Quantification of number of filopodia per cell. Data was pooled from 3 independent experiments. Number of cells analyzed: control: n = 40 cells, WT MRTF-A OX n = 60 cells, ΔSAP MRTF-A OX n = 43 cells, ΔSRF MRTF-A OX n = 58 cells). **E)** Quantification of filopodia length. Data was pooled from 3 independent experiments. Number of filopodia analyzed: control: n = 495 filopodia, WT MRTF-A OX n = 1373 filopodia, ΔSAP MRTF-A OX n = 986 filopodia, ΔSRF MRTF-A OX n = 933 filopodia. For all quantification, the ROUT method (Q = 1%) was used to remove outliers. One-way ANOVA was used to compare means. **** indicates p value < 0.0001, *** indicates p value < 0.001, ** indicates p value< 0.01, and * indicates p value < 0.05.

### mDia2 is a mediator of MRTF-A induced promotion of breast cancer cell motility

Membrane protrusion is orchestrated by the coordinated actions of a wide range of actin-binding proteins (ABPs). These ABPs include proteins involved in maintenance of the monomeric actin pool via G-actin sequestering, nucleotide exchange on G-actin, nucleation and/or elongation of actin filaments, disassembly of actin filaments, filament severing, capping and crosslinking of actin filaments. Therefore, to determine which of these major classes of ABPs are transcriptionally altered upon various functional perturbations of MRTF-A, we performed bulk RNA-sequencing analyses of both 2D and 3D cultures of control vs various MRTF-A overexpressing sublines of MDA-231 cells (GSE253047). We then derived differentially expressed genes (DEGs) based on the absolute Log_2_fold change (FC) (>1.5) in gene expression, false-discovery rate (FDR) <0.001, and the expression level cut-off set to 0.1 TPM (transcripts per million). Heat-maps of the DEGs between the cell lines (**Fig S5A**) and principal component analyses (PCA) (**Fig S5B**) revealed that overexpression of ΔSRF-MRTF-A had the most dramatic impact on the gene expression profile in MDA-231 cells. The transcriptional profiles of major ABPs belonging to the different functional categories in 2D and 3D cultures of each of the four sublines are shown in the form of heat-plots in **Fig 5A** and **Fig 5B**, respectively. We specifically filtered for genes which were differentially expressed (defined as greater than 2 Z-scores - Z scores were calculated using the mean and standard deviation of the control cells for each gene) between WT-MRTF-A and ΔSRF-MRTF-A overexpressers, and in the opposite directions relative to control cells mirroring the migration pattern of these groups of cells (**Fig 5C**). From this list, we identified *DIAPH3* to be the only gene that was reproducibly altered at an mRNA level in a consistent direction (i.e. elevated and downregulated upon overexpression of WT-MRTF-A and ΔSRF-MRTF-A, respectively) in both 2D and 3D cultures.

**Figure 5.**
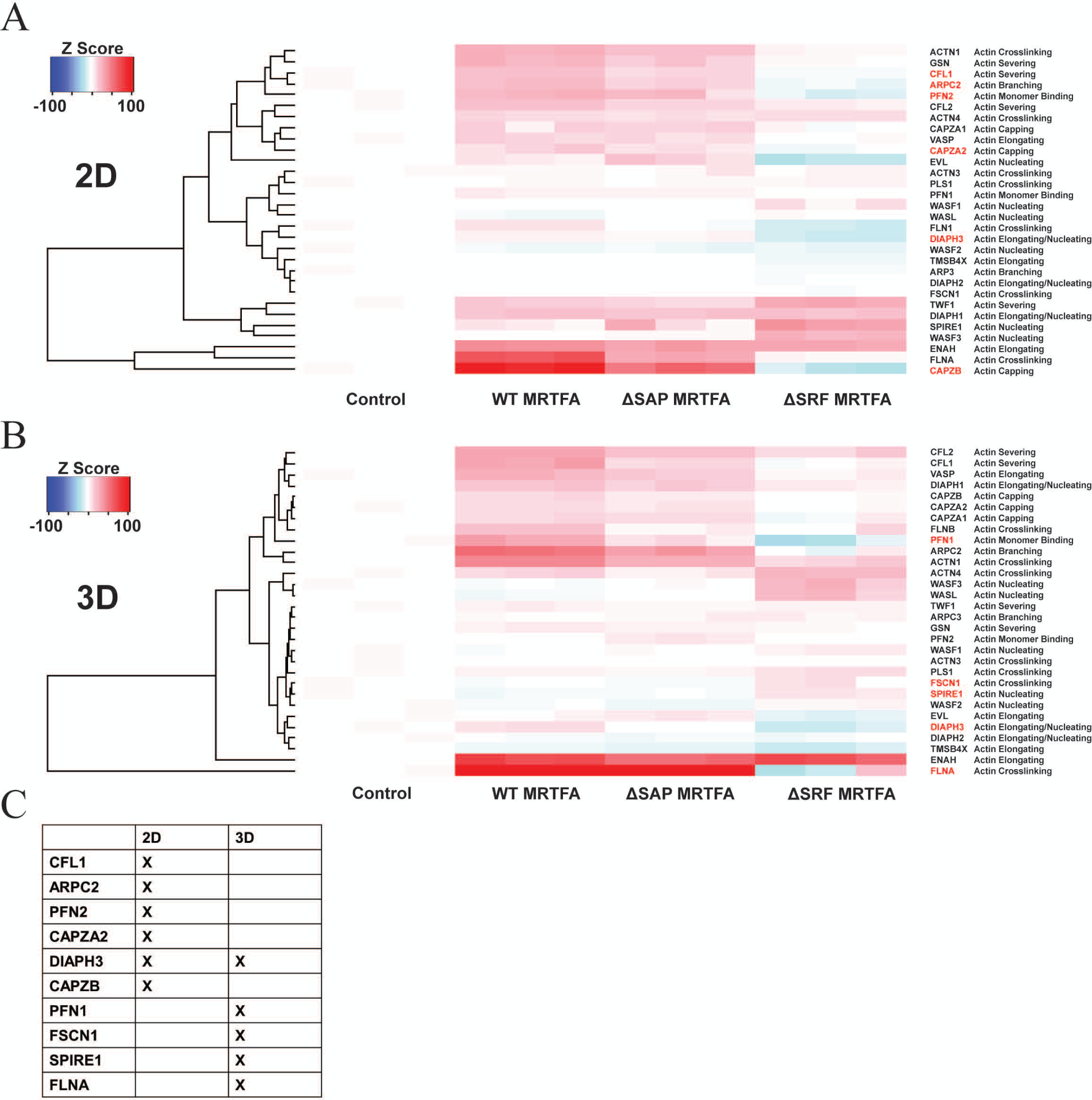
MRTF-dependent changes in the expression of ABPs in 2D and 3D culture environments. A-B) Heat-plots showing MRTF-A-dependent changes in the mRNA expression (represented as Z-scores based on the control group’s standard deviation and mean) of major classes of ABPs in MDA-231 cells based on RNA-sequencing data of cells cultured on 2D tissue culture (*panel A*) and 3D matrigel-on-top (*panel B*) substrates with 3 biological replicates. **C)** Table summarizing the list of those ABP-encoding genes which differed in expression by at least 2 Z-scores from the control group, and in the opposite directions in WT- vs ΔSRF-MRTF-A overexpression group.

Formin-family gene *DIAPH3* (encodes mDia2) is one of the three human orthologs of drosophila *Diaphanous* (Schoen, et al. 2013). Like other members of formin-family proteins, mDia2, when activated through its direct interaction with Rho GTPase (Rif1 specifically in the case of mDia2), promotes nucleation and elongation of actin filaments and formation of F-actin-based structures (Gorelik, et al. 2011; Stastna, et al. 2012). Although mDia2 is most well-known for its ability to induce filopodia formation (Schirenbeck, et al. 2005; Stastna, et al. 2012), overexpression of constitutively active mDia2 also leads to longer actin filaments in lamellipodia and improved lamellipodial protrusion in motile cells (Yang, et al. 2007). Given mDia2’s well-established role in actin polymerization and membrane protrusion, we further investigated whether mDia2 could be a potential mediator of the MRTF-dependent regulation of cell migration. First, to verify MRTF-A dependent regulation of mDia2 expression at the protein level, we performed immunoblot analyses of MDA-231 cell lysates which showed that mDia2 expression is elevated upon overexpression of either WT- or ΔSAP-MRTF-A but severely downregulated when ΔSRF-MRTF-A is overexpressed **(Fig 6A)**. Note that although WT- and ΔSAP-MRTF-A overexpression-induced mDia2 upregulation was a consistent feature across all experiments, it did not reach statistical significance because of the variability in fold-change of upregulation. Consistent with these overexpression results, co-silencing of MRTF isoforms, either transiently in MDA-231 cells or stably in D2A1 (a murine breast cancer cell line) cells, downregulated mDia2 expression (**Figs 6B-C**). However, the expression of mDia1 (another member of formin family of proteins that is encoded by the *DIAPH1* gene, closely related to and has overlapping functions with mDia2) either remained unaffected or was minimally diminished (not statistically significant) in these cell lines (**Figs 6B-C**), suggesting that MRTF has a specific effect on mDia2 expression without having a global effect on the expression of all formins.

**Figure 6.**
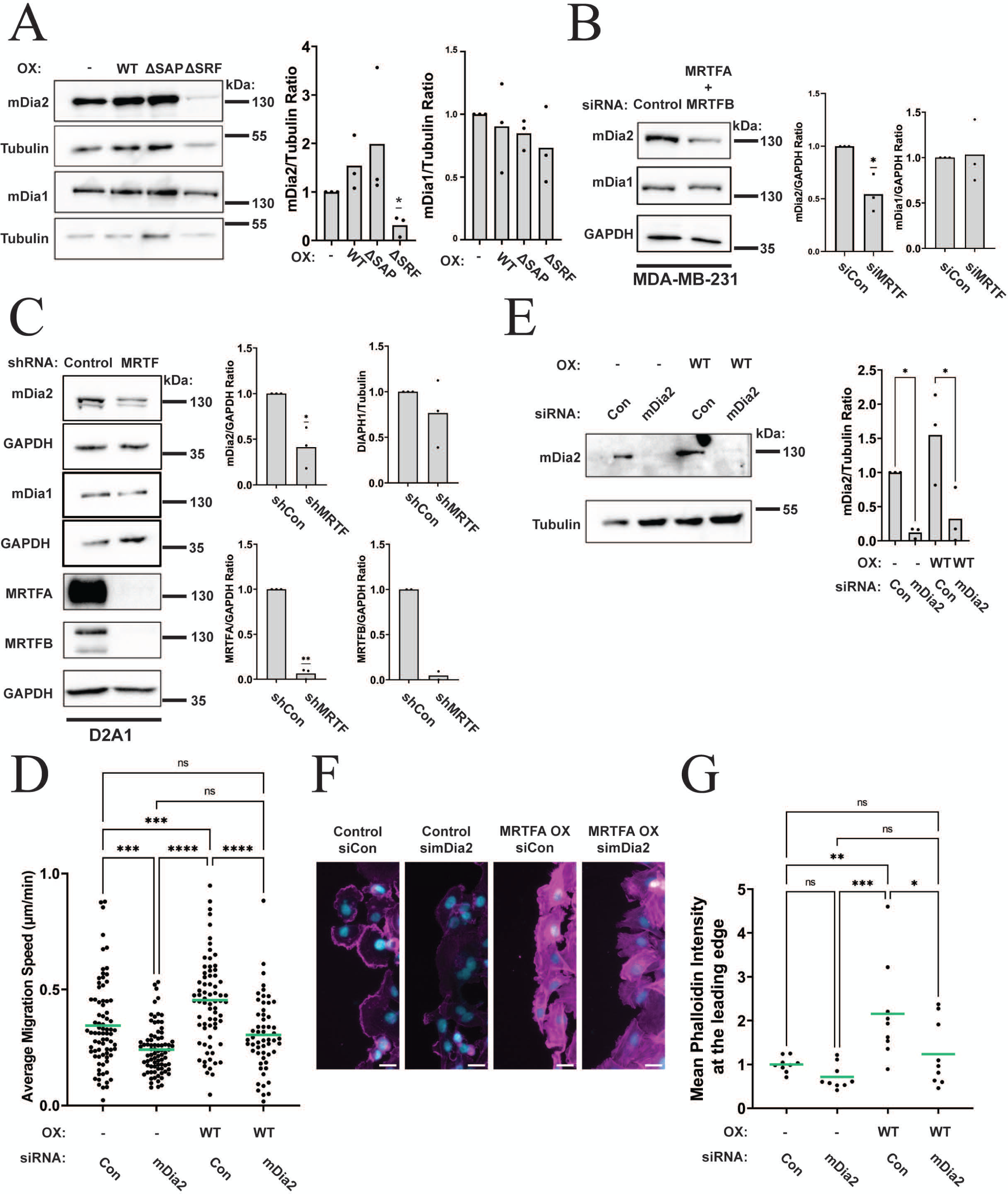
Demonstration of mDia2 is a mediator of MRTF-A induced promotion of cell motility and actin polymerization at the leading edge. A) Representative immunoblots and the associated quantification showing the relative expression levels of mDia1 and mDia2 after 48h dox treatment in the indicated cell lines (OX – overexpression). **B-C)** Representative immunoblots and the associated quantification showing the changes in the expression levels of mDia1 and mDia2 in MDA-231 (*panel B*) and D2A1 (*panel C*) after transient and stable knockdown of MRTF isoforms, respectively. GAPDH and tubulin blots serve as the loading controls. Immunoblot data were quantified from 3 experiments, and one-sample T-test was used to determine significance. **D-E)** Dot blots (with indicated means) in *panel D* represent the average migration speed of control vs MRTF-A overexpressing MDA-231 cells 72 hrs after siRNA-mediated knockdown of mDia2 in 2D motility assays. As control, each group of cells was transfected with non-targeting control (con) siRNA. Dox-induced MRTF-A overexpression was turned on for 48 hrs before cells were subjected to siRNA transfection. Number of cells pooled from 3 independent experiments analyzed in each group: control (si-Con) – 78, control (si-mDia2) – 79, WT-MRTF-A OX (si-Con) – 73, and WT-MRTF-A OX (si-mDia2) – 56. Immunoblot in *panel E* demonstrates mDia2 knockdown (tubulin blot – loading control). **F-G)** Representative phalloidin staining images (*panel F;* scale bar = 10µm) at the migrating front of a wound-healing assay and quantification of mean phalloidin intensity at the leading edge of cells (*panel G*) (images were acquired 3 hours after the monolayer was scratched, and 9 leading fronts were analyzed/group). Data points are pooled from 3 independent experiments with 3 technical replicates per experiment. For all quantification, the ROUT method (Q = 1%) was used to remove outliers. One-way ANOVA was used to compare means. **** indicates p value < 0.0001, *** indicates p value < 0.001, ** indicates p value< 0.01, and * indicates p value < 0.05.

Next, we analyzed 2D random migration of control vs WT-MRTF-A overexpressing sublines of MDA-231 with or without forced knockdown of mDia2 (non-targeting control siRNA transfectants served as control). As expected, overexpressing WT-MRTF-A increased the average migration speed of MDA-231 cells by ∼32%. Knockdown of mDia2 in control and WT-MRTF-A overexpressing cells decreased average migration speed compared to their control siRNA treated counterparts by 30% and 33%, respectively, mitigating the differences in average migration speed between the control and MRTF-A overexpression groups of cells (**Fig 6D**; immunoblot shown in **Fig 6E** demonstrates mDia2 knockdown). These results suggest that MRTF-A stimulates cell migration in an mDia2-dependent manner. To further examine a potential role of mDia2 in MRTF-A-induced changes in actin polymerization at the leading edge, we pre-established monolayers of control and MRTF-A overexpressing cells with or without forced knockdown of mDia2 and evaluated mDia2-induced changes in actin polymerization at the lamellipodial edge in polarized migrating cells in a wound-healing assay. In parallel, we examined mDia2’s involvement in MRTF-induced changes in global F-actin content in a culture setting that did not involve wound healing. As expected, MRTF-A overexpression increased the overall F-content in cells (marked by promotion of actin stress fibers) and actin polymerization at the leading edge (**Figs 6F** and **Fig S6**). Knockdown of mDia2 expression alone was not sufficient to block the MRTF-A-induced increase in overall F-actin content (**Fig S6**). However, MRTF-A-induced increase in actin polymerization at the leading edge was dramatically reversed upon mDia2 depletion (**Figs 6F-G**). These data indicate that mDia2 is a mediator of MRTF-A driven stimulation of actin polymerization at the leading edge and cell migration.

Given our observation of a drastic loss of mDia2 expression upon overexpression of ΔSRF-MRTF-A, we next asked to what extent SRF is responsible for regulating cellular mDia2 expression. To address this question, we performed transient silencing of SRF expression in MDA-231 cells. As per these experiments, SRF silencing led to a moderate reduction in mDia2 expression although the effect did not reach statistical significance (**Fig 7A)**. Since mDia2 downregulation upon loss of SRF expression was not nearly as robust as elicited by either MRTF depletion or overexpression of ΔSRF-MRTF-A, we speculated that there might be additional mechanisms underlying MRTF-dependent regulation of mDia2 that does not directly involve SRF’s action. It has been shown that MRTF-A/SRF and YAP/TEAD transcriptional signaling complexes crosstalk and lead to activation of each other through their ability to control cytoskeletal dynamics in fibroblasts (Foster, et al. 2017). Another study also found MRTF-B’s ability to promote YAP/TEAD activity in MDA-231 cells does not require its SRF’s interaction (Kim, et al. 2017). We found that silencing YAP expression substantially reduces mDia2 expression in MDA-231 cells (**Fig 7B**). This is consistent with ChIP-seq data of binding the YAP-TEAD4 transcriptional complex to the promoter of *DIAPH3* gene (Zanconato, et al. 2015). Interestingly, our immunofluorescence studies revealed that neither the total expression nor the nuclear-to-cytoplasmic ratio of YAP in MDA-231 cells is impacted by overexpression of either WT-MRTF-A or ΔSRF-MRTF-A (**Figs 7C-E**). In light of these findings, we speculate that MRTF-A could have both SRF-dependent and -independent mechanisms of regulating mDia2 expression, the latter through priming YAP-TEAD activity or possibly through additional YAP- and SRF-independent mechanisms (as schematized in **Fig 7F**). However, more in-depth studies are needed to fully elucidate the mechanisms.

**Figure 7.**
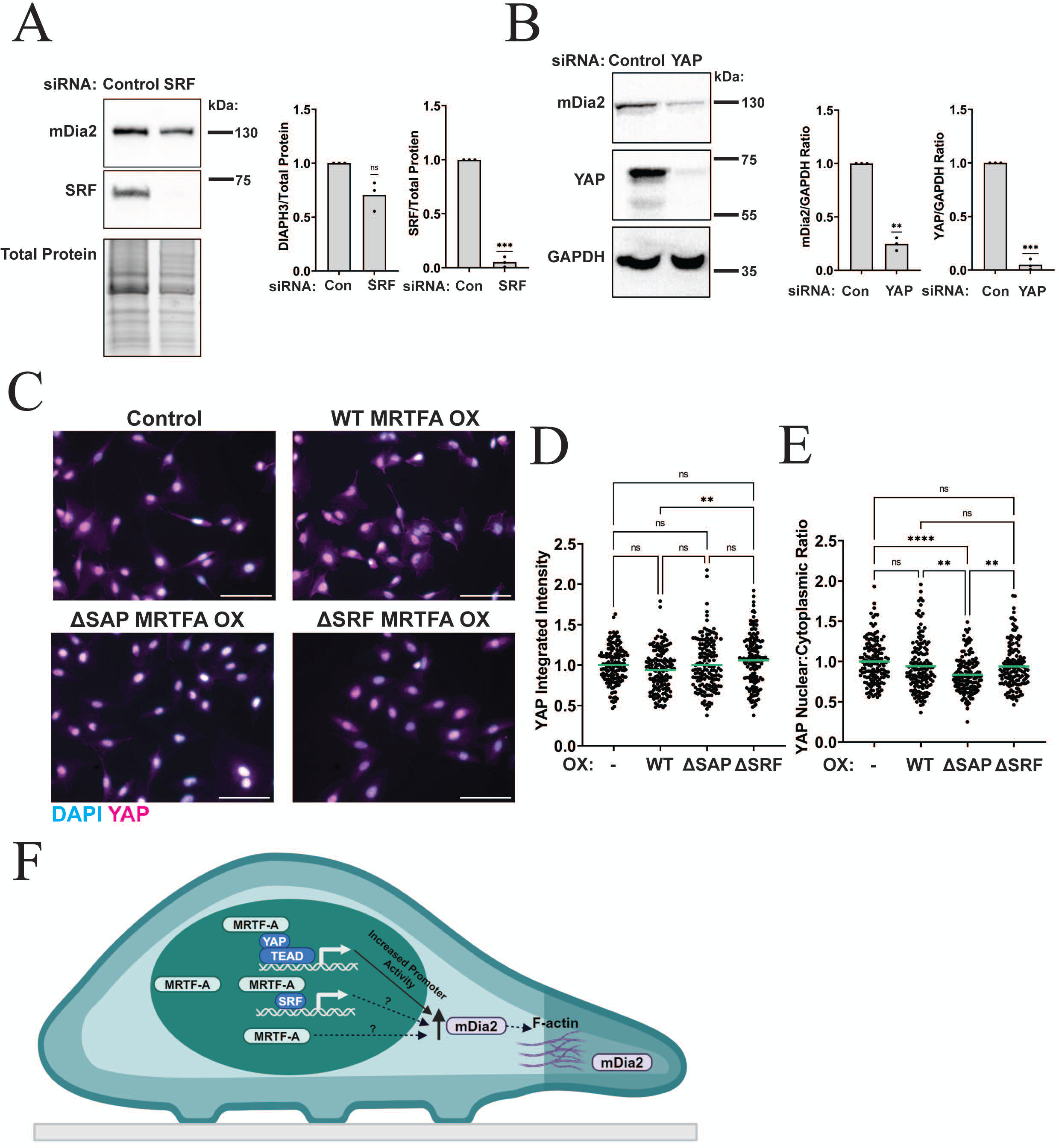
Potential mechanisms of MRTF-dependent regulation of mDia2 expression. A) Representative immunoblots and associated quantification (n=3 experiments) of changes in the expression of mDia2 and SRF in MDA-231 cells 96 hrs after transfection with SRF-specific vs control-siRNA (total lysate band quantification was used as a normalizing factor). **B)** Representative immunoblots and associated quantification (n=3 experiments) of changes in the expression of mDia2 and YAP in MDA-231 cells 96 hrs after transfection with YAP-specific vs control-siRNA (n=3 experiments; GAPDH blot serves as a loading control). One-sample T-test was used to determine significance for panels A and B. **C-E)** Representative immunostaining images of YAP (*panel C*; with DAPI counterstain) and quantification of the total YAP intensity (*panel D*) and the nuclear-to-cytoplasmic intensity ratio of YAP in the indicated groups of cells (*panel E*) pooled from three independent experiments (OX – overexpression). Number of cells analyzed: Control n = 145 cells, WT MRTF-A OX n = 150 cells, ΔSAP MRTF-A OX n = 150 cells, and ΔSRF MRTF-A OX n = 151 cells. For all quantification, the ROUT method (Q = 1%) was used to remove outliers. One-way ANOVA was used to compare means. **** indicates p value < 0.0001, *** indicates p value < 0.001, ** indicates p value< 0.01, and * indicates p value < 0.05. **F**) A proposed schematic model of MRTF-dependent regulation of mDia2 expression and stimulation of cell migration utilizing mDia2-mediated actin polymerization at the leading edge.

### Increased MRTF activity positively correlates with local invasion and distant metastases in human breast cancer

Next, we performed multiplexed quantitative immunohistochemistry (IHC) of a human BC tissue microarray (TMA) representing 361 clinical samples (clinico-pathological features of the TMA are detailed in the “Experimental Procedures” section), staining for MRTF-A, MRTF-B, pan-cytokeratin (CK: to distinguish CK+ tumor cells from CK-stromal cells by image segmentation), and nuclear marker DAPI (representative images are shown in **Figs 8A, D**). Pan-CK staining was used to segment the TMA spots into tumor and stromal regions (the tumor masks are shown in **Figs 8B** and **8E)**. Since local invasion into the stroma and/or lymph node infiltration are characteristic features of malignant cells, we performed two metrics of analyses to query a relationship between tumor-cell intrinsic MRTF activity in an isoform-specific manner and malignancy potential of tumor cells. First, we compared the expression of each MRTF isoform between CK+ cancer cells in the bulk tumor vs those dispersed in the stromal region. The underlying assumption is that the latter category of cells represents locally invading cancer cells enabling their separation from the bulk tumor. Interestingly, as per these analyses, the total expression level of MRTF-A was significantly lower in cancer cells dispersed in the stroma than those in the bulk tumor (**Fig 8G**); MRTF-B expression in cancer cells was not statistically different between the two compartments (**Fig 8J**). However, when we analyzed the nuclear-to-cytoplasmic (N/C) content of each MRTF isoform (as a surrogate indicator of the transcriptionally active form of MRTF), the average N/C ratios of both MRTF-A and MRTF-B in CK+ cells were significantly higher in the stromal *vs.* the bulk tumor regions (**Figs 8H, K)**. Representative images of MRTF-A and MRTF-B nuclear localization in cancer cells in the stromal regions (marked by asterisks) and cancer cells in the bulk tumor (outlined with a dotted line) are shown in **Figs 8C** and **8F**. Second, we performed correlation analyses between the N/C ratio of each MRTF isoform with the number of involved lymph nodes in the clinical specimens. When we performed these analyses including all tumor samples, we did not see any significant correlation between the N/C ratio of either of the two MRTF isoforms and the number of lymph node involvement **(Fig S7)**. However, when these analyses were performed in a molecular subtype-specific manner, we found a significant positive correlation between the N/C ratio of MRTF-A (but not MRTF-B) and the number of lymph node involvement exclusively in lymph node positive TNBC samples (**Figs 8I, L**). In summary, these TMA data provide an initial indication for higher MRTF activity positively correlated with malignant traits of tumor cells in human BC, more prominently in TNBC.

**Figure 8.**
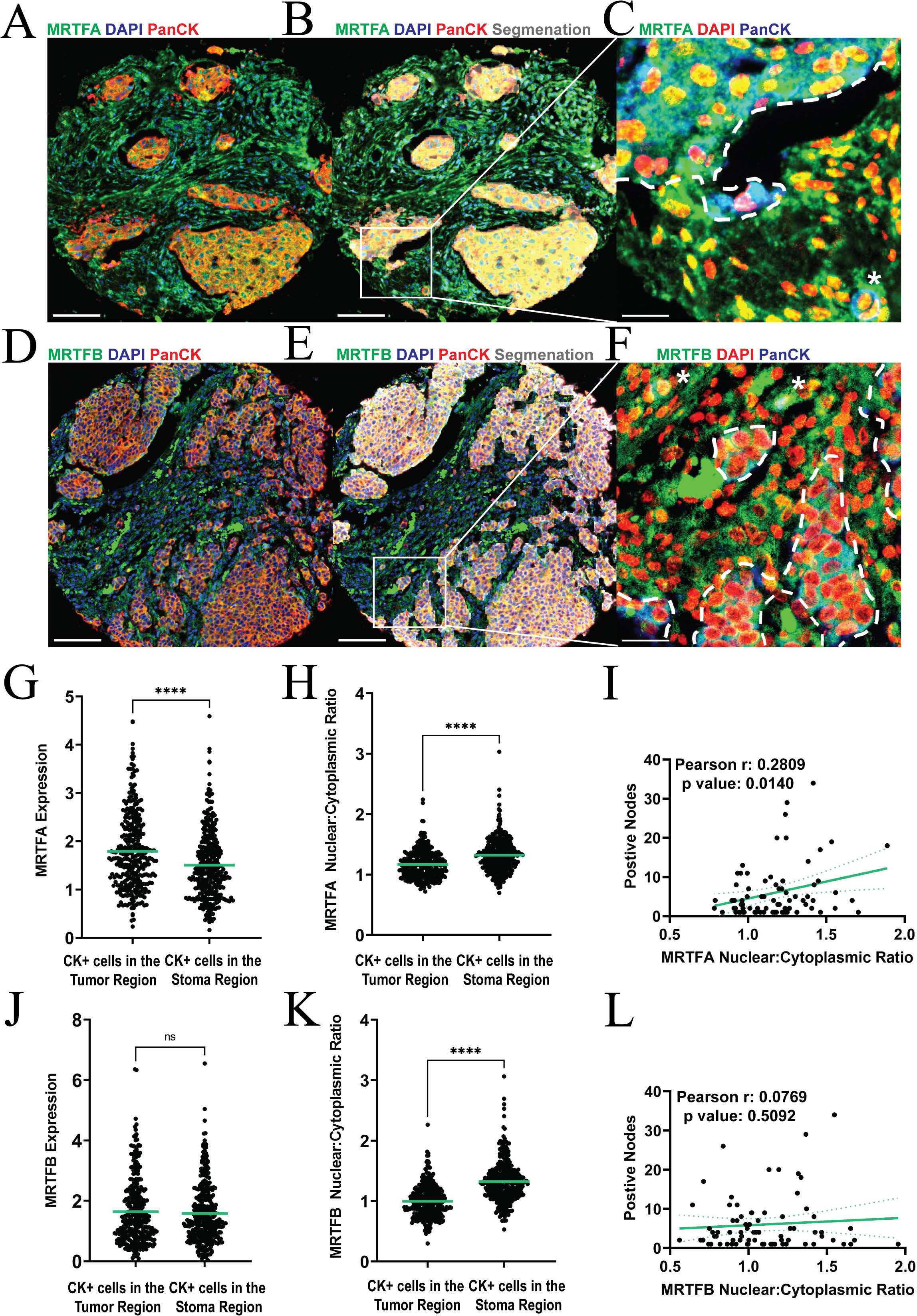
Multiplexed quantitative IHC-based assessment of association between subcellular localization of MRTF isoforms and malignancy-associated features in human BC. A, D) Representative images of a breast cancer TMA spot showing the tumor microenvironment stained for various markers including MRTF-A, MRTF-B, pan-cytokeratin (Pan-CK), and DAPI. Scale bar = 100µm **B, E)** Representative images of the segmentation of TMA spots into tumor and stromal regions. Tumor regions are overlayed with grey projection. **C, F)** Zoomed-in regions of TMA from the indicated regions of panel B and E showing expression and sub-cellular localization of MRTF-A and MRTF-B isoforms in cells, respectively. Pan-CK+ cells were identified as cancer cells. The dotted white line indicates the tumor regions and the white asterisk marks Pan-CK+ cells in the stroma. Scale bar = 25µm. **G, J)** Quantification of the average MRTF-A (*panel G*) and -B (*panel J*) expression of Pan-CK+ cells in the segmented tumor and stromal regions. (p value < 0.0001, n = 339 samples). **H, K)** Quantification of the average nuclear-to-cytoplasmic (N/C) ratio of MRTF-A (*panel H*) and -B (*panel K*) in Pan-CK+ cells in the segmented tumor and stromal regions (p < 0.0001, n = 339 samples). The N/C ratio was determined by dividing the mean MRTF intensity in the nuclear area by the mean MRTF intensity in the cytoplasmic area in an isoform-specific manner. **I, L)** Correlation plots showing the association between the number of positive lymph nodes and N/C ratio of the individual MRTF isoform (*panel I*: MRTF-A; *panel L* – MRTF-B) in Pan-CK+ cells collectively from the tumor and stroma compartments in TNBC (the histology grading scores for HER2, ER, and PR of these tumor samples were determined to be either 0 or 1).

Although our TMA studies utilized the nuclear localization status of MRTF as a surrogate marker for the active form of MRTF, we recognize that it is still only an approximate indicator of MRTF activity. This is because the transcriptional activity of MRTF can be also regulated by nuclear actin (Miralles, et al. 2003) and various post-translational modifications (Panayiotou, et al. 2016; Holstein, et al. 2020). Furthermore, since MRTF has both SRF-dependent/SAP-independent and SRF-independent/SAP-dependent transcriptional activities (Gurbuz, et al. 2014), nuclear localization status alone fails to provide any insight into different transcriptional modalities of MRTF. Therefore, we sought to investigate which specific transcriptional signature of MRTF is predominantly associated with malignancy traits in human breast cancer. Given that our TMA studies provide stronger evidence for MRTF-A’s association with the malignant traits than for MRTF-B in TNBC, we overlapped DEGs between 2D cultures of control and WT-MRTF-A overexpressing MDA-231 cells obtained from our studies with published DEGs of 2D cultures of MDA-231 cells in response to stable co-silencing of both MRTF isoforms restricted to identical FDR and FC criteria as previously reported by the Treisman group (Medjkane, et al. 2009). From these analyses, we identified genes that were transcriptionally altered in the opposite directions in knockdown vs overexpression setting to serve as a high-stringency MRTF-A gene signature. Function-specific gene signatures were then developed by first removing DEGs which were represented in the WT-but not the corresponding functional-mutant MRTF-A overexpression setting. Then the gene signature score (GSS) of downregulated genes were subtracted from GSS of upregulated genes as illustrated in the Venn diagrams (**Fig 9A**) and the table (**Fig 9B**) to generate scores which account for both positive and negative gene regulation by MRTF-A’s function. Further details of the generation of the gene signature and the genes which composed these signatures can be found in the “Experimental Procedures” and **Table S1**. Note that since the WT-MRTF-A overexpression induced increase in mDia2 transcript level was ∼22% in 2D culture (compared to 66% increase in 3D culture), it did not meet the criterion for gene signature construction and is therefore not represented in **Table S1**. Next, we ran gene signature variance analyses (GSVA) using our generated gene signatures on primary breast tumors and pair-matched metastases harvested from various anatomical sites (ovary, gastrointestinal tract, brain, and bone) of BC patients (Zhu, et al. 2019; Cosgrove, et al. 2022). Although there were site specific differences in the trend, in the case of ovary and brain metastases, there was a trend of higher MRTF-A SRF specific GSS (9 out of 12 in the ovary, and 17 out of 22 in the brain). (**Fig 9C**). While site-specific analyses did not reach statistical significance (except for brain metastases), likely because of small size for each anatomical site or possible site-specific MRTF signaling requirements for metastasis, pooled analyses of all soft tissue metastases revealed a statistically significant enrichment of SRF function-specific but not SAP-dependent gene signature of MRTF-A in metastases relative to the primary tumor (**Fig 9D)**. Collectively, these data are indicative of higher MRTF-SRF activity associated with greater metastatic potential of tumor cells in human BC.

**Figure 9.**
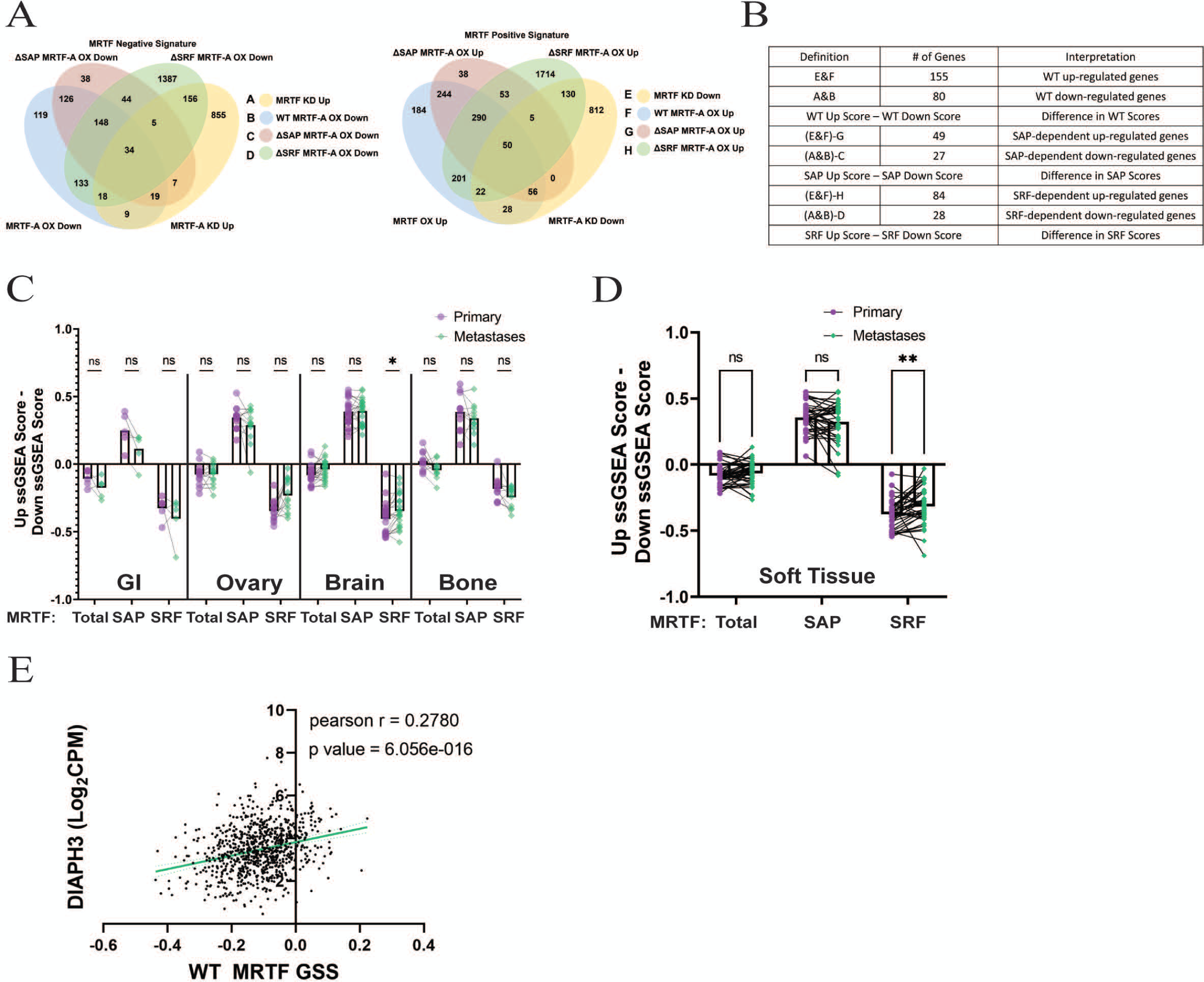
Evidence for elevated MRTF-SRF gene signature in distant soft-tissue metastases relative to pair-matched primary tumors in human BC. A) Venn Diagrams representing the overlap of DEGs between the indicated groups to identify genes that are negatively (left) and positively (right) regulated by MRTF-A (cutoffs: absolute Log_2_fold change > 1.5, false-discovery rate (FDR) <0.001, and expression level cut-off =0.1 TPM). **B)** Table showing the composition of the calculated gene signature scores constructed from the overlapping Venn diagrams. Differential gene signatures capturing both positive and negative functions of MRTF-A were generated by subtracting the positive gene signature scores from the negative gene signature scores as indicated. **C-D)** Comparison of MRTF gene signature scores (GSS) in metastases *vs* pair-matched primary tumors (panel C - individual metastatic site, and panel D – all soft tissue sites (GI tract, ovary and brain) combined). A non-parametric Wilcoxon-rank sum test with false discovery rate correction was used to compare the median GSS between groups (the adjusted p-values are indicated above the plots).

Finally, to investigate a potential connection between MRTF, mDia2 and malignancy in human BC, we queried publicly available The Cancer Genome Atlas (TCGA) transcriptome dataset. Consistent with our cell line data, TCGA data analyses revealed a moderate positive correlation between MRTF-A GSS and mDia2/*DIAPH3* mRNA expression (**Fig 9E**). Note that a moderate correlation was not totally unexpected since a) mDia2 expression might be regulated by other transcription factors besides MRTF, and b) whole tumor (contains contribution from other cell types in the tumor microenvironment) transcriptome data was analyzed in our studies. We also performed an exploratory analysis of mDia2/*DIAPH3* mRNA expression in pair-matched primary tumors vs distant metastasis. Although we saw a trend of elevated mDia2 expression in 30 out of 51 distance metastases (7 out of 11 in bone, 14 out of 22 in brain, 5 out of 13 in ovary, and 4 out of 5 gastrointestinal tract) relative to their pair-matched primary tumors (**Fig S8**), unfortunately, the small sample size precludes us from reaching a definitive conclusion on mDia2’s association with malignancy in human BC.

## DISCUSSION

In the present study, we report several novel findings. First, while a number of previous studies showed defects in migration and invasion of cells in developmental and pathological (cancer) contexts upon loss of function of either MRTF or SRF (Han, et al. 2004; Somogyi and Rorth 2004; Medjkane, et al. 2009; Evelyn, et al. 2010; Franco, et al. 2013; Weinl, et al. 2013; Hayashi, et al. 2014; Costello, et al. 2015; Record, et al. 2015; Watanabe, et al. 2015; Gau, et al. 2017), to the best of our knowledge this study provides the first direct experimental evidence for MRTF’s ability to promote cell migration and invasion through its interaction with SRF. Second, this study establishes a role of mDia2 in MRTF-dependent regulation of actin polymerization at the leading edge and cell migration. Third, this is the first study that correlates subcellular localization of MRTF in an isoform-specific manner with the clinical features of malignancy and compares function-specific gene signature of MRTF between metastases harvested from different anatomical sites and matched primary tumors to collectively demonstrate a positive association between MRTF activity and metastatic potential of tumor cells in human BC.

Our studies demonstrate that MRTF’s ability to promote both 2D and 3D cell migration requires SRF’s interaction, while the SAP-domain function of MRTF is selectively important for 3D cell migration. Previously another group investigated the effect of overexpression of WT and the identical mutants of MRTF-A on transwell migration of HC11 mammary epithelial cells (Gurbuz, et al. 2014). Our results are somewhat at odds with their finding of increased chemotactic migration of HC11 cells upon overexpression of ΔSRF-MRTF-A and defect in migration induced by overexpression of ΔSRF-MRTF-A. While their findings, particularly the one related to ΔSRF-MRTF-A’s effect on cell migration, are hard to reconcile with the literature, we suspect there could be several reasons that could potentially explain the discrepancy in the results. First, a key difference is the use of cell line model - while MDA-231 cells are mesenchymal (lacks E-cadherin expression), HC11 cells are epithelial in nature where there could be a confounding factor of the effect of disruption of MRTF-SRF interaction on cell-cell interaction. Second, chemotactic migration of HC11 cells was assessed in a long-term assay (22 hrs) which can have a confounding effect of cell proliferation. Third, transwell migration is also facilitated by deformability (important for cell migration through the pores) and gradient-sensing ability of cells. It is possible that weakening of actin cytoskeleton induced by ΔSRF-MRTF-A could improve the deformability of cells enabling them more adaptable for migration through the transwell pores.

Although our results from short-term (12 hr) single-cell invasion and long-term (96 hr) tumor spheroid invasion assays appear contradictory in terms of the effect of WT MRTF-A overexpression on cell invasion, we believe that our single cell invasion data scored by time-lapse imaging more accurately reflects how MRTF elevation promotes the intrinsic invasiveness of BC cells. There are several possible explanations underlying this apparent discrepancy. First, a lower directed invasion readout of tumor spheroid in MRTF-A overexpression setting could be partly due to reduced directional persistence of movement of these cells in a 3D environment as revealed by our single-cell time-lapse imaging data. Second, we previously reported that WT-MRTF-A overexpression increases the proliferation accelerating the primary tumor growth of MDA-231 cells when implanted in mice (Gau, et al. 2022). Both proliferation and migration are energy-dependent processes. According to “Go-or-Grow” hypothesis, highly proliferating tumor cells undergo phenotypic switching to reduce migratory ability as a way of channeling energy utilization toward proliferation (Giese, Kluwe, et al. 1996; Giese, Loo, et al. 1996; Giese, et al. 2003), which may partly explain the reduced invasiveness of WT-MRTF-A overexpressers relative to control cells in a long-term assay. Third, MMP (matrix metalloproteinase)-mediated ECM proteolysis is an important feature of tumor cell invasion, and it is possible that perturbation of MRTF can modulate the expression of certain MMPs and tissue inhibitors of MMPs (TIMPs) to remodel the ECM architecture, additionally impacting the mode (ameboid vs mesenchymal) of 3D cell migration. Nevertheless, our tumor spheroid invasion results suggest that excessively high MRTF activity may also not be optimal for directed ECM invasion of BC cells.

An interesting general finding in our cell migration/invasion studies is that overexpression of MRTF-A mutants in some cases not only reversed the hypermigratory phenotype of WT-MRTF-A overexpressing cells but also actively inhibited cell migration. There could be two possible scenarios underlying this observation. One possibility is that overexpression of a specific MRTF-A mutant has a dominant negative action suppressing the endogenous MRTF activity (this is conceptually not impossible since MRTF isoforms homo- and hetero-dimerize). Alternatively, the mutant forms of MRTF have additional unknown gain- or loss-of functions which partially contribute to phenotypic changes. When we probed our RNAseq data, we found that several known MRTF-SRF target genes (example: ACTA2, TAGLN, CTGF) are transcriptionally upregulated upon overexpression of WT-MRTF-A as expected. Meanwhile, ΔSRF-MRTF-A overexpression reversed the increased expression of these genes, but not below the control which would be expected for a dominant negative. Given that gene expression is often regulated by a complex and redundant network of transcriptional pathways, the dominant negative aspect cannot be formally ruled out. However, given the pattern of gene expression of MRTF-SRF target observed in the RNA-seq data, the latter scenario is a clear possibility. It is also possible that inactivation of one function of MRTF-A may hyperactivate another function.

In this study, we demonstrate that MRTF promotes mDia2 expression and cell migration in an mDia2-dependent manner. Although mDia2 expression is substantially diminished at the protein level when MRTF expression is silenced and even more so, when ΔSRF-MRTF-A is overexpressed, silencing SRF expression only modestly reduced mDia2 level. One possible interpretation of these findings is that MRTF has both SRF-dependent and -independent regulation of mDia2. As per the ENCODE project data, *DIAPH1* (encodes mDia1) but not *DIAPH3* (encodes mDia2) is a bona-fide transcriptional target of SRF (Consortium 2004). A previous study found that mRNA-binding protein IGF2BP3 increases mDia2 mRNA stability involving m^6^A-modification of the transcripts (Wu, et al. 2022). As per our RNA-seq data, we saw >2-fold increase in IGF2BP3 mRNA expression in MDA-231 cells upon overexpression of WT-MRTF-A. Therefore, IGF2BP-mediated mRNA stability control could be one of the mechanisms of MRTF-A-dependent upregulation of mDia2 expression. Additionally, our findings raise the possibility of YAP’s involvement as another arm of MRTF’s regulation of mDia2. We saw that overexpression of either WT- or SRF-binding deficient mutant of MRTF-A has no effect on nuclear localization of YAP, a finding that resonates with a previous finding that MRTF-B potentiates the activation of YAP/TEAD complex through recruitment of additional gene regulatory factors (e.g. NCOA3) without requiring SRF’s interaction (Kim, et al. 2017). Therefore, it will be interesting to see in the future whether a similar pathway is involved in SRF-independent regulation of mDia2 downstream of MRTF activity. Mass-spectrometry analyses of immunoprecipitated WT- vs ΔSRF-MRTF-A might offer additional mechanistic insight into gain- and loss-of-functions of MRTF-A in terms of its ability to bind to other ligands when its interaction with SRF is prevented.

Several cancer-related studies have previously documented a role of mDia2 in modulating tumor cell aggressiveness albeit in a context- and study-specific manner. For example, pan-cancer analyses revealed that mDia2 is upregulated in multiple types of human cancer (breast, cervical, head and neck squamous carcinoma, glioblastoma and several others) and serves as a prognostic manner in certain cancers (Chen, et al. 2023). The expression of mDia2 is upregulated in osteosarcoma, and silencing mDia2 expression suppresses proliferation, migration/invasion and metastasis of osteosarcoma cells (Zhang, et al. 2021). In the context of hepatocellular carcinoma (HCC), while one study documented mDia2’s ability to stimulate migration/invasion and metastatic capability of HCC cells (Dong, et al. 2018), another study found a non-coding RNA-mediated exon skipping leading to reduced expression of mDia2, further linking to epithelial-mesenchymal transition, migration, invasion and metastasis of HCC cells (Su, et al. 2023). Conflicting results have also been reported in the literature with regard to mDia2’s role in BC. In one study, silencing mDia2 induced ameboid migration promoting invasion and metastasis of oncogenic HRAS-transformed human mammary epithelial cells (Hager, et al. 2012). Depletion of mDia2 also induces nuclear shape instability, promoting the migratory ability of breast and prostate cancer cells (Reis-Sobreiro, et al. 2018). Contrasting with these findings, another study found that all diaphanous proteins including mDia2 are required for invadopodia formation and efficient ECM invasion of MDA-231 breast cancer cells (Lizarraga, et al. 2009). Our findings that mDia2 silencing reduces actin polymerization at the lamellipodial edge and reduces migration of MDA-231 cells are consistent with previously demonstrated mDia2’s role in lamellipodial protrusion (Yang, et al. 2007). Although we are not unable to reconcile our findings with anti-migratory effect of mDia2 in BC cells as shown in some of the previous studies (Hager, et al. 2012; Reis-Sobreiro, et al. 2018), we speculate study-specific differences in model systems and experimental assays may partly underlie the dichotomy in the findings.

Only a handful of studies to date have assessed MRTF expression/activity in human BC. One study found higher mRNA expression of MRTF-A and select MRTF-A target genes (ACTA2/α-smooth-muscle actin, SM22/TAGLN, CD49e/integrin-α5, and CCN2/CTGF) in the primary tumor associated with shorter overall survival of a small set of luminal (n=129) and mixed (n=104) breast cancer patient cohorts (Seifert and Posern 2017). Another study revealed MRTF-A’s SRF-independent/SAP-dependent gene expression as a prognostic signature of highly proliferative BC with adverse clinical outcome (Gurbuz, et al. 2014). However, these studies involved gene expression analyses of bulk tumor, which does not distinguish the effect of MRTF in tumor cell from stromal cell populations. We also recognize that the overall survival data is influenced by various aspects of tumor cell behavior (growth, dissemination, and survival) in addition to other patient-specific confounding variables that are unrelated to cancer. Therefore, we currently lack an explicit understanding of whether tumor-cell-intrinsic expression/sub-cellular localization of any particular isoform of MRTF is correlated specifically with malignancy-associated features in human BC. Our TMA IHC data showed that cancer cells infiltrated in the stromal region have higher N/C ratio of both MRTF-A and MRTF-B relative to those present in the bulk tumor. A possible interpretation of this finding could be that invading subpopulation of tumor cells upregulate MRTF activity. Interestingly, N/C ratio of only MRTF-A (but not MRTF-B) in cancer cells exhibited a positive correlation with lymph node infiltration in TNBC. The underlying reason for isoform-specific selectivity of MRTF for association with lymph node infiltration in TNBC is not clear. However, since lymph node infiltration also involves intravasation, it is possible there might be an isoform-specific effect of MRTF activity in cancer cells on endothelial barrier disruption (a key aspect of intravasation) which can be investigated in the future. Encouragingly, consistent with our cell line-based experimental data showing critical importance of MRTF/SRF interaction for tumor cell migration and invasion, we provide clinical evidence for elevated SRF-specific gene signature of MRTF in distant metastases relative to matched primary tumor in human disease. However, we acknowledge that it is impossible to rule out that MRTF/SRF’s requirement for metastatic outgrowth of breast cancer cells (Gau, et al. 2022) is an additional contributing reason for elevated MRTF/SRF activity in distant metastases.

## MATERIALS AND METHODS

### Cell culture and transfection

Generation and culture of GFP/luciferase-expressing sublines of MDA-231 expressing various dox-inducible MRTF-A constructs have been recently reported by our group (Gau, et al. 2022). D2A1 cell line (originally provided to us by Dr. William Schiemann, Case Western Reserve University) were transduced with lentivirus encoding either MRTF-A/B shRNA (Vectorbuilder vector ID - VB200825-1246dwj) or non-targeting control shRNA (Vectorbuilder vector ID - VB151023-10034), and selected/maintained with puromycin in DMEM media supplemented with FBS (Corning, catalog# MT35011CV) and antibiotics (100 U/ml penicillin and 100 μg/ml streptomycin; Thermo Fisher, catalog# 15070063). HEK-293 cells were cultured in DMEM-F12 media supplemented with FBS (Corning, catalog# MT35011CV) and antibiotics (100 U/ml penicillin and 100 μg/ml streptomycin; Thermo Fisher, catalog# 15070063). For transient gene silencing, siRNAs were transfected using Lipofectamine RNAiMAX (Thermo Fisher, catalog #13778100). The following siRNA were used at a concentration of 2.5 nM (MRTF-A, MRTF-B, and YAP) or 5 nM (mDia2): custom single target 3’ UTR MRTF-A (Dharmacon – Sequence: 5’-GUGUCUUGGUGUAGUGUAAUU-3’), Smart-Pool MRTF-B siRNA (Santa Cruz, catalog #sc-61074), Smart-Pool mDia2 siRNA (Dharmacon Catalog ID: E-018997-00-0005), Smart-Pool YAP1 siRNA (Dharmacon Catalog ID: L-012200-00-0010). For transient plasmid transfection of HEK-293 cells, X-tremeGENE™ HP DNA Transfection Reagent (Roche Life Science Products; Catalog# XTGHP-RO) was used in a 1 µl X-tremeGene: 1 µg of plasmid ratio.

### Immunoblotting

Total lysate was extracted in Pierce^TM^ IP Lysis buffer (Thermo Scientific – Catalog #87787) containing 0.3% SDS. Prior to loading samples in the acrylamide gel the samples were boiled in 1X sample buffer. For immunoblotting, the following antibodies at the following concentrations were used: polyclonal anti-MRTF-A antibody (Cell Signaling – 14760S; 1:1000), polyclonal anti-MRTF-B antibody (abcam – ab235101; 1:1000), polyclonal anti-GAPDH antibody (Sigma Aldrich – ABS16; 1:1000), monoclonal anti-mCherry antibody (Cell Signaling – E5D8F; 1:1000), polyclonal anti-DIAPH3 (Abcam – ab245660; 1:1000), polyclonal DIAPH1 antibody (Abcam – ab11173; 1:1000), monoclonal Flag antibody (Sigma Aldrich – B3111; 1:1000), monoclonal SRF antibody (Active Motif – 61385; 1:2000), and monoclonal YAP (Cell Signaling – 14074; 1:1000).

### GST-pulldown Experiment

BL21(DE3) E. Coli cells (New England Biolabs – Catalog #C2527I) were transformed with pGEX4T-SRF plasmid (Addgene #121100 – kindly deposited by Paul Herring) and GST-SRF expression was induced by overnight IPTG (0.1 mM) treatment at room temperature. GST-SRF was purified from bacterial lysate through immobilization on glutathione-linked agarose beads (Thermo Scientific – Catalog#16100) and washed several times before incubating GST-SRF-conjugated bead with lysates of HEK-293 cells transiently transfected with flag-tagged constructs encoding expressing either WT or K237A/K238A/H239A mutant of MRTF-A (Joy, et al. 2017) for 1 hour at 4°C.

### Time-lapse cell motility assay

For 2D motility assays, cells were sparsely plated on collagen coated (10 µg/ml) 24-well plate overnight and imaged for 3h at 10 min intervals with a 10x objective. For 3D motility assays, cells were first seeded on top of 100 µl polymerized 1 mg/ml Collagen type-1 matrix in a 96 well plate. After 4 hours of incubation at 37C to allow for cell adhesion, an additional 100 µl of 1 mg/ml Collagen type-1 was added on-top of the cells. This ensured the cells were completely immersed in the collagen-1 matrix, but in the same plane to allow for convenient imaging. The cells were then imaged for 12h at 30 min intervals with a 10x objective. For both 2D and 3D motility assays, the onstage environmental chamber was maintained at 37°C and 5% CO_2_ for the entire duration of time-lapse imaging. Multiple beacons for imaging were selected for frames with single cells in each well. Cells were segmented and tracked using Ilastik (Berg, et al. 2019) and trajectories were constructed using centroid coordinates of cell areas. Only cells that were present in the imaging frame for greater than 2 hours were counted. Persistence, shown as standard deviation of Δθ, was calculated using Δθ with θ_i, i+1_ = cos^−1^(x_i+1_ − x_i_)/sqrt ((x_i+1_ − x_i_)^2^) + ((y_i+1_ − y_i_)^2^) where x and y are the position coordinates. Directionality was calculated by calculating the end-to-end distance divided by total distance traveled per cell. For experiments involving dox-inducible overexpression, cells were treated with dox for 48h prior to imaging. For experiments involving gene knockdown, time-lapse imaging studies were performed 72h after siRNA transfection.

### Spheroid invasion assay

Spheroids composed of 3000 uninduced (i.e., non-doxycycline treated) cells were formed for 48 hours in 10 µg/ml Collagen-1 (Gibco - Catalog #A10483-01) in the wells of a round-bottom, low-adhesion 96-well plates (Corning – product number 7007). After 48 hours of formation, 1 µg/ml doxycycline was added to the wells for 24 hours. Collagen-1 (prepared at 2 mg/ml) was added to the media in the wells to a final concentration of 1 mg/ml and allowed to polymerize for 1 hour prior to the addition of serum-containing media supplemented with 1 µg/ml dox. After allowing cells to invade for 96 hours, the spheroids were fixed in 3.7% formaldehyde for 1h, permeabilized overnight in 0.5% Triton-X at 4°C and stained with DAPI and Acta-Stain 670 phalloidin (Cytoskeleton – catalog # PHDN1) overnight at 4°C. Spheroids were imaged at 0h and 96h with a 2.5X objective. Spheroid invasion data were pooled from 7 independent experiments with each experiment having 5 technical replicates; However, the spheroids which were not able to be imaged (primarily due to not being in the center of the round bottom plate) were excluded from the study.

### Kymography analysis

Time-lapse imaging of randomly migrating cells was performed over 10 minutes at a 2-second time interval with a 20X objective. Kymographs were constructed using compiled slices of the time-lapse frames along multiple 1-pixel lines drawn across the protruding edge of a cell using ImageJ. Kymographs were then processed in ImageJ by background subtraction followed by automated local thresholding about the mean intensity. Protrusions were identified from the binary thresholded images using R. Local maxima and minima were identified in the kymographs using the following method. First, local regression (LOESS) was performed to generate a smoothed curve (plotted as a black line in **Fig 4A**) using the coordinates of the thresholded leading edge. Next, on a rolling basis the highest point in the span of 7 time points (12s) was assigned to each point along the curve. This generated a curve, herein referred to as the highest local point loess regression curve (plotted in green in **Fig 4A**). Finally, the loess regression curve was subtracted from the highest point loess regression curve. The points at which the results of the subtracted curves were equal to zero were treated as the local maxima. The same algorithm was used to identify local minima. Local maxima less than 2 pixels high were excluded from the analysis. The frequency of protrusion, net protrusion distance and duration of protrusion ascent were then calculated based on these identified local minima and maxima. Kymographs were generated from three independent experiments with 3 technical replicates (3 wells were plated) per experiment. Cells from multiple wells were selected from analysis across the two experiments. ∼2 kymographs were constructed from each cell analyzed.

### F-Actin staining

Cells cultured on the wells of a collagen-1-coated (10 µg/ml) 24-well plates were washed with PBS, fixed with 3.7% formaldehyde for 15 minutes, permeabilized with 0.5% Triton-X for 5 minutes, and stained with Acta-Stain 670 phalloidin for 30 minutes (Cytoskeleton – catalog # PHDN1) at room temperature. Counterstaining for nuclei with DAPI was subsequently performed for 1 minute at room temperature. Fluorescence images acquired with a 20X objective were processed in ImageJ for background subtraction and automatic thresholding before quantifying the integrated intensity on a per cell basis by utilizing the “Analyze Particles” function. For wound-healing assay, cells were seeded in the wells a 24-well plate at 80% confluency and allowed to reach confluency overnight. A wound was made in each well using a 200 µl pipet tip, and cells were allowed to migrate for 3 hours before performing phalloidin staining and fluorescence image acquisition of the leading edge as described above. Images thresholded for phalloidin-positive region were exported to R for calculation of the average fluorescence intensity at the 5 pixels across the entire leading edge. The wound healing assay was conducted across three independent experiments with three technical replicates each (three wells were plated). One image was taken per well.

To induce filopodia, sparsely plated MDA-231 cells were serum-starved overnight and stimulated with 1 μM bradykinin (R&D Systems; Catalog #3004) for 30 minutes before fixation and staining with phalloidin as described above. Automated image quantification was performed using FiloQuant, an automated filopodia detection plug-in for ImageJ (Jacquemet 2023).

### Immunostaining

Cells cultured on the wells of a collagen-1-coated (10 µg/ml) 24-well plates were washed with PBS, fixed with 3.7% formaldehyde for 15 minutes, permeabilized with 0.5% Triton-X for 5 minutes, blocked with 10% goat serum for 1 hour, incubated with 1:100 dilution of primary antibodies for YAP (Cell Signaling – 14074) or MRTF (Cell Signaling – 14760S) overnight at 4C, washed with PBS, and incubated with 1:100 dilution of Cy5 Goat Anti-Rabbit IgG (Jackson labs; Catalog #111-175-144) for 1h at room temperature. Following PBS washes, the cells were stained with DAPI.

### Multiplexed quantitative IHC

Breast cancer TMA was purchased from Yale University (YTMA 49-13) and has been used in many previous studies for cancer biomarker identification and validation (Zerkowski, et al. 2007; Giltnane, et al. 2009; Chakraborty, et al. 2018). The TMA was composed of spots from 647 breast cancer patient samples. After quality assessment post-staining, 360 of the spots were deemed quality and used for further evaluation. 18.6% of the samples were stage 1, 37.5% of the samples were stage 2, 26.9% of the samples were stage 3, 6.1% of the samples were stage 4, and 10.8% of the samples did not have a determined stage. 47 % of samples were ER+ and 53% of samples were ER- as determined by the histology score (a score of 0 or 1 was counted as ER-). The tissue microarray was specifically procured to have approximately equal numbers of ER- and ER+ samples. Automated staining of TMA was performed on the Leica Bond RX. For staining, Akoya Bioscience’s Opal 6-Plex Detection Kit (cat# NEL871001KT) was used according to the manufacturer’s instructions. The antibodies used in the staining process are as follows: MRTF-A (abcam – ab113264 – 1:200), MRTF-B (abcam – ab235101 – 1:500), and pan-cytokeratin (Clone AE1/AE3 – Santa Cruz – sc81714 – 1:150). Akoya Bioscience’s PhenoimagerHT platform was used for whole slide scanning. Akoya Biosciences InForm® analysis software was used for spectral unmixing, tissue/cell segmentation, and phenotyping. The minimum size threshold to be counted as a tumor region was 2000 pixels. Any single or clusters of pan-CK+ cells less than 2000 pixels were counted as cancer cells present in the stroma compartment.

### RNA-sequencing and gene signature construction

For RNA sequencing, MDA-231 cells were cultured on either 2D tissue culture dishes or on top of 3D matrigel substrate (as per protocol of matrigel-on-top culture detailed in (Gau, et al. 2022), and induced with dox for 72h for transgene (WT vs mutant forms of MRTF-A) expression prior to RNA-collection with the Qiagen RNeasy Mini Plus Kit following manufacturer’s protocol. Samples were submitted to the University of Pittsburgh HSCRF Genomics Research Core for RNA-sequencing. The core facility prepared the RNA library using TruSeq Stranded Total Ligation (Ribo-Zero Plus) protocol (Illumina) and performed paired-end reads on the Illumina NextSeq 2000 platform. Samples were sequenced to a minimum of 22 million reads per sample. Quality of sequencing reads was determined with FastQC before transcripts from the Ensembl v.93 transcriptome (hg38) were quantified with Salmon v0.14.0. Gene-level expression was calculated using tximport v.1.12.0, differential gene expression was calculated with DESeq2 v.1.24.0, and BiomaRt v.2.46.0 was used to annotate gene names. Genes with a minimum expression value greater than 1 transcripts per million (TPM) in either control or experimental group, an adjusted *p*-value less than 0.01, and an absolute Log_2_fold change greater than 1.5 were considered differentially expressed. Raw read data and processed data can be accessed from an SRA/GEO submission that is currently in progress. Procedures of gene signature generation are detailed in the “Results” section. GSVA package was used to calculate the ssGSEA scores (Hanzelmann, et al. 2013).

### Transcriptome analyses of paired primary and metastatic breast tumors

The dataset consists of 51 pairs of primary and paired metastases from cohort in University of Pittsburgh. DNA/RNA was extracted from formalin-fixed paraffin-embedded (FFPE) tissue from patient-matched primary breast tumors and resected metastases using the Qiagen GeneRead DNA FFPE kit using standard protocols. Sample quality and concentration were assessed by Qubit and fragment analysis. RNA library was prepared with TruSeq Stranded Total RNA (Degraded RNA) v2 RNA Exome Library and TS RNA Access capture protocol (Illumina), and RNA expression passing quality control was quantified with Salmon and normalized in log-normalized counts before ssGSEA calculation. Detailed description of data processing is as described in (Cosgrove, et al. 2022)

### Statistics and data representation

All statistical tests were performed in GraphPad Prism 9 software. Specifics of statistical tests are indicated in the figure legends. Data were plotted in GraphPad Prism 9 or in R (ggplot2).

## ACKNOWLDEGMENTS

IE was supported by NIBIB T32-EB001026 and NCATS TL1-TR001858. VY was supported by a summer research internship from the School of Engineering at the University of Pittsburgh. DG was supported by a National Cancer Center fellowship, NCI K99-CA267180 and T32-HL129964. Work in the Roy lab was supported by grants from NIH (R01CA248873, R01CA271095, and R21EY032632). Fangyuan Chen was a former visiting research scholar at the University of Pittsburgh School of Medicine supported by funds from The China Scholarship Council and Tsinghua University.

## AUTHOR CONTRIBUTIONS

IE designed experiments, performed experiments, analyzed data, and wrote manuscript; VY and DG performed experiments and analyzed data, MJ performed experiments, analyzed data and wrote manuscript, DB and FC performed data analyses and wrote manuscript; JA and PC performed experiments, AVL and PL oversaw some aspects of data analyses, PR conceived the overall study design, supervised the project, oversaw experiments, edited/wrote manuscript and acquired funding.

