## Supplementary Info for "mDia2 is an important mediator of MRTF-A-dependent regulation of breast cancer cell migration"

Eder, I. et al. Fig S1

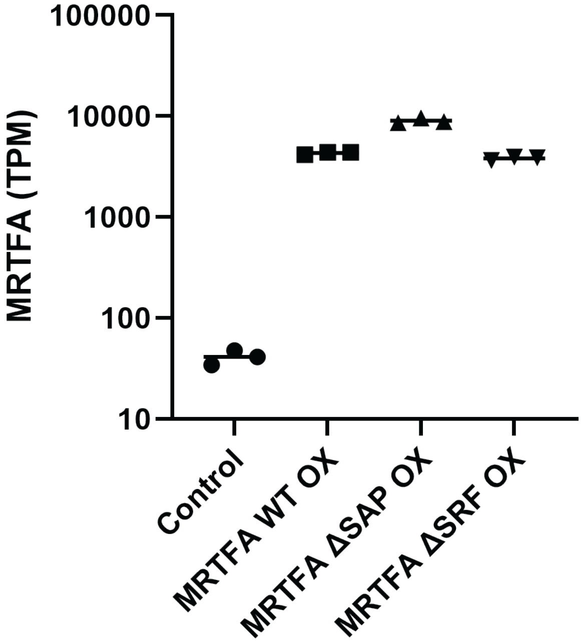

**Figure S1.** Relative mRNA expression of MRTF-A (in TPM) in the indicated sublines of MDA-231 cells in 2D culture.

Eder, I. et al. Fig S2

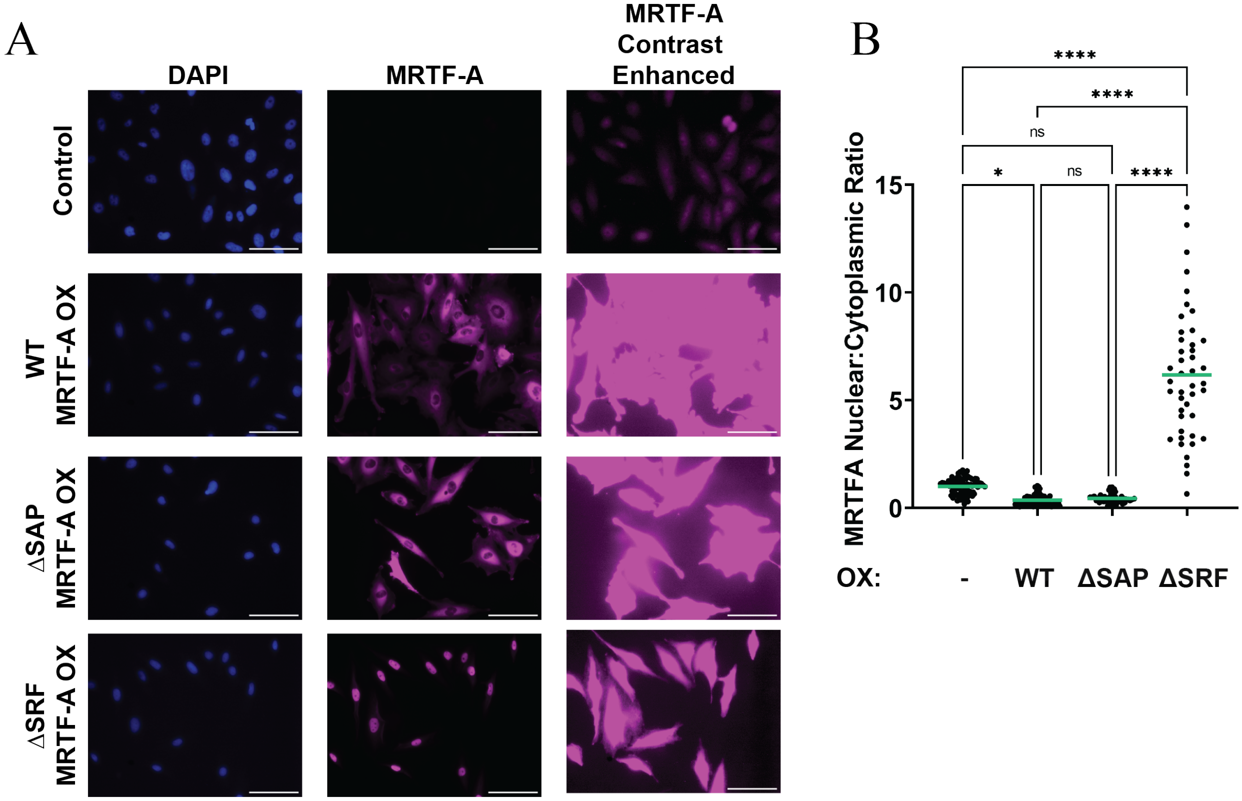

**Figure S2. Sub-cellular localization of WT vs mutant MRTF-A. A)** Representative MRTF-A immunostaining (with DAPI counterstaining) images of the indicated groups of MDA-231 cells at normal (used for quantification) and high contrast (shown here to display endogenous MRTF-A staining) OX – overexpression. **B)** Quantification of the nuclear-to-cytoplasmic ratio of MRTF-A in the indicated cell lines. Cells analyzed per condition: Control, n= 84 cells. WT MRTF-A OX, n = 75 cells, ΔSAP MRTF-A OX, n = 44 cells, and ΔSRF MRTF-A OX, n = 46 cells. One-way ANOVA was used to compare means. **** indicates p value $< 0.0001$, * indicates p value $< 0.05$

Eder, I. et al. Fig S3

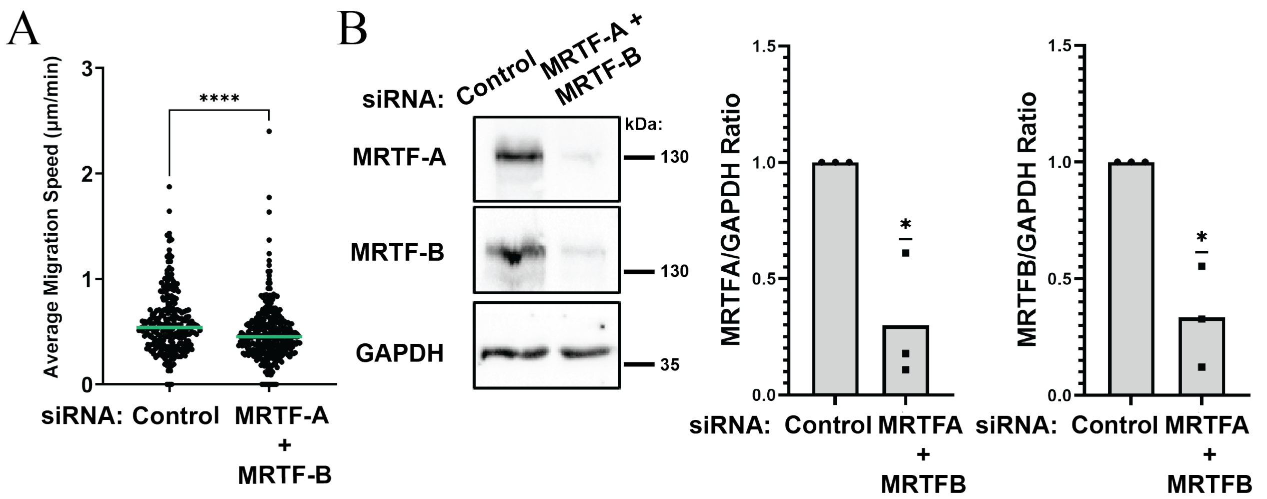

**Figure S3. Effect of transient co-silencing of MRTF isoforms on 2D random motility of MDA-231 cells. A)** Dot plot representing the average migration speed of individual MDA-231 cells 72 hrs after transfection with either non-targeting siRNA or pooled siRNAs targeting MRTF-A and MRTF-B isoforms. Cells were pooled across three independent experiments (siControl: n = 262; siMRTF-A/B: n =313). A student's T-test was used to compare the means. **** indicates p value < 0.0001. **B)** Immunoblot-based validation of MRTF-A and -B knockdown in MDA-231 cells (GAPDH serves as the loading control). A one sample T-test was used to compare the means of the experimental group to the control group. * indicates p < 0.05.

Eder, I. et al. Fig S4

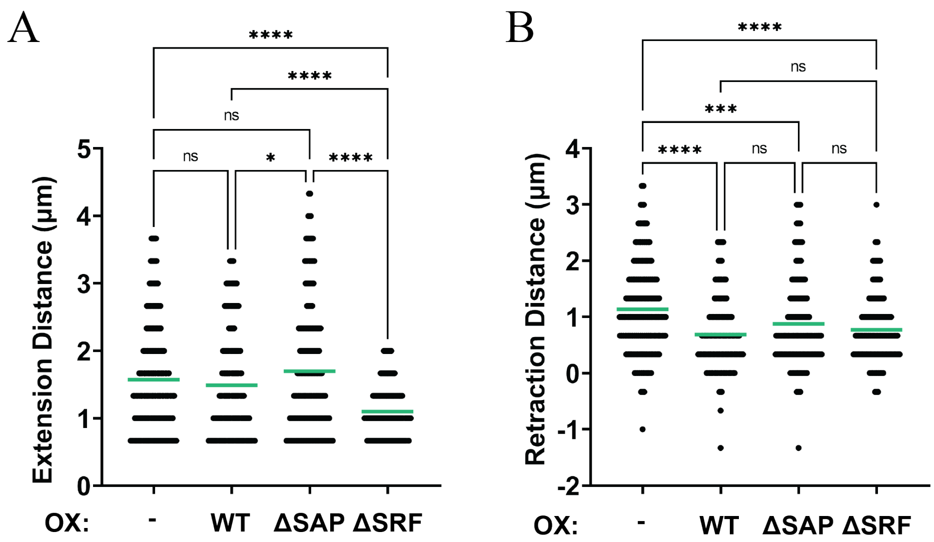

**Figure S4.** Quantification of the extension (*panel A*) and retraction (*panel B*) of each individual membrane protrusion per indicated group (OX – overexpression). Number of protrusions analyzed: control - 400, WT-MRTF-A OX –229, ΔSAP-MRTFA OX -174, and ΔSRF-MRTFA OX - 169 protrusions. For all quantifications, outliers were removed using the ROUT method (Q = 1%). One-way ANOVA was performed. **** indicates p value < 0.0001, *** indicates p value < 0.001, ** indicates p value< 0.01, and * indicates p value < 0.05.

Eder, I. et al. Fig S5

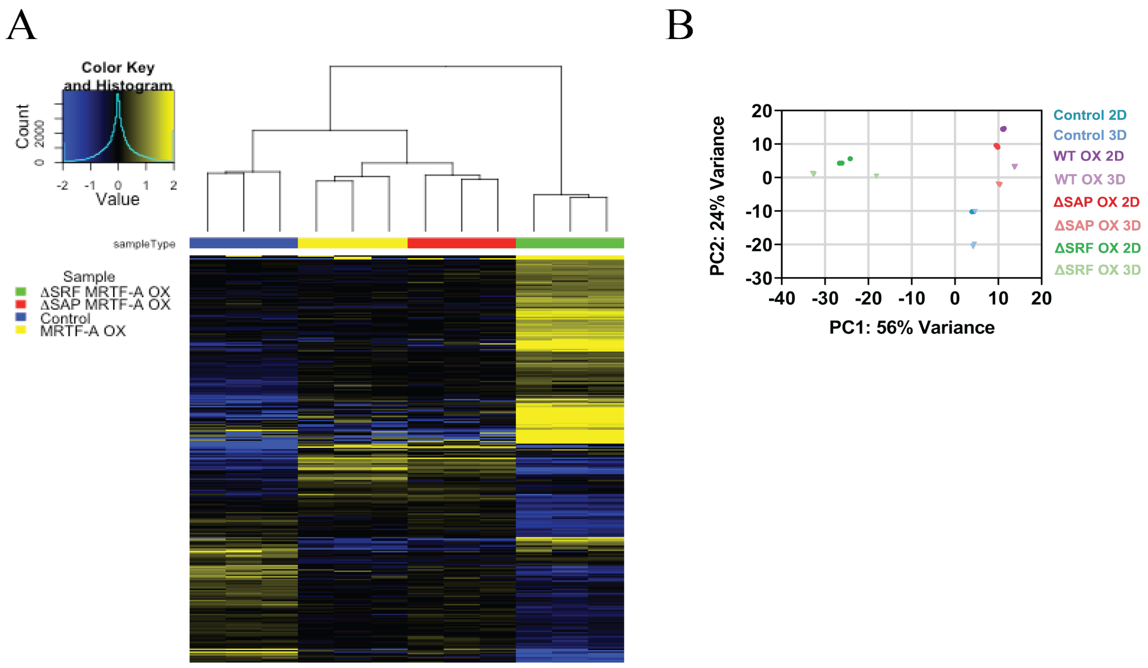

**Figure S5. Changes in the global transcriptomic profile of MDA-231 cells induced by overexpression of WT- vs mutant forms of MRTF-A. A)** Heat-plot showing the expression of differentially expressed genes (DEGs) (cut off: $\log_2$fold change > 1.5, TPM > 1, FDR < 0.001) compared to the control. **B)** Principal component analysis demonstrating most robust changes in transcriptome induced by overexpression of ΔSRF-MRTFA OX.

Eder, I. et al. Fig S6

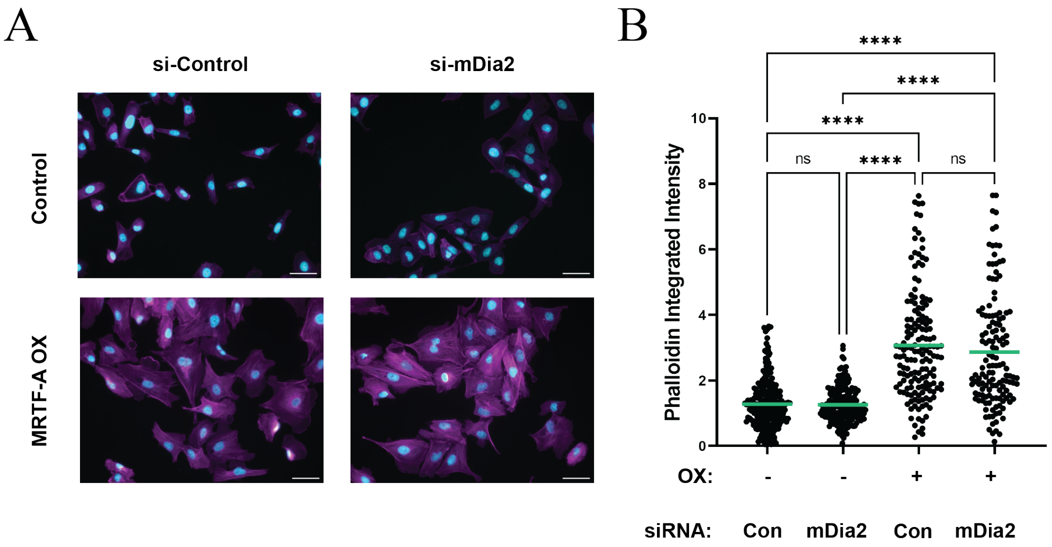

**Figure S6. Effect of mDia2 silencing on MRTF-A induced changes in bulk F-actin polymerization. A)** Representative 20X images of Phalloidin and DAPI stained control and WT MRTF-A overexpressing cells treated with non-targeting siRNA or mDia2 siRNA. Scale bar = 20μm. **B)** Quantification of the integrated intensity of phalloidin per cell. Cells were pooled from 3 independent experiments. (OX – overexpression). Control/siCon: n = 96 cells, Control/si-mDia2: n = 115 cells, WT-MRTF-A OX/siCon: n = 77 cells, WT-MRTF-A OX/si-mDia2: n = 89 cells. One-way ANOVA was used to compare means. **** indicates p value < 0.0001, and ** indicates p value < 0.001.

Eder, I. et al. Fig S7

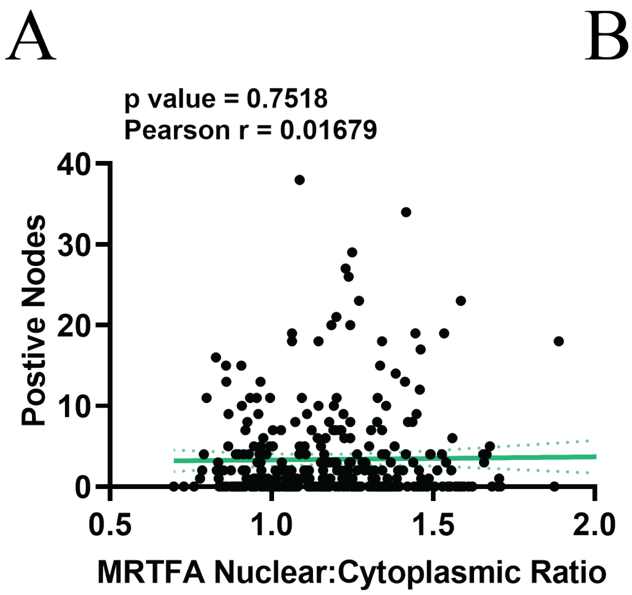

**Figure S7.** Correlation plots showing the association between the number of positive lymph nodes and N/C ratio of the individual MRTF isoform (*panel A*: MRTF-A; *panel B* – MRTF-B) in Pan-CK+ cells collectively from the tumor and stroma compartments in all samples.

Eder, I. et al. Fig S8

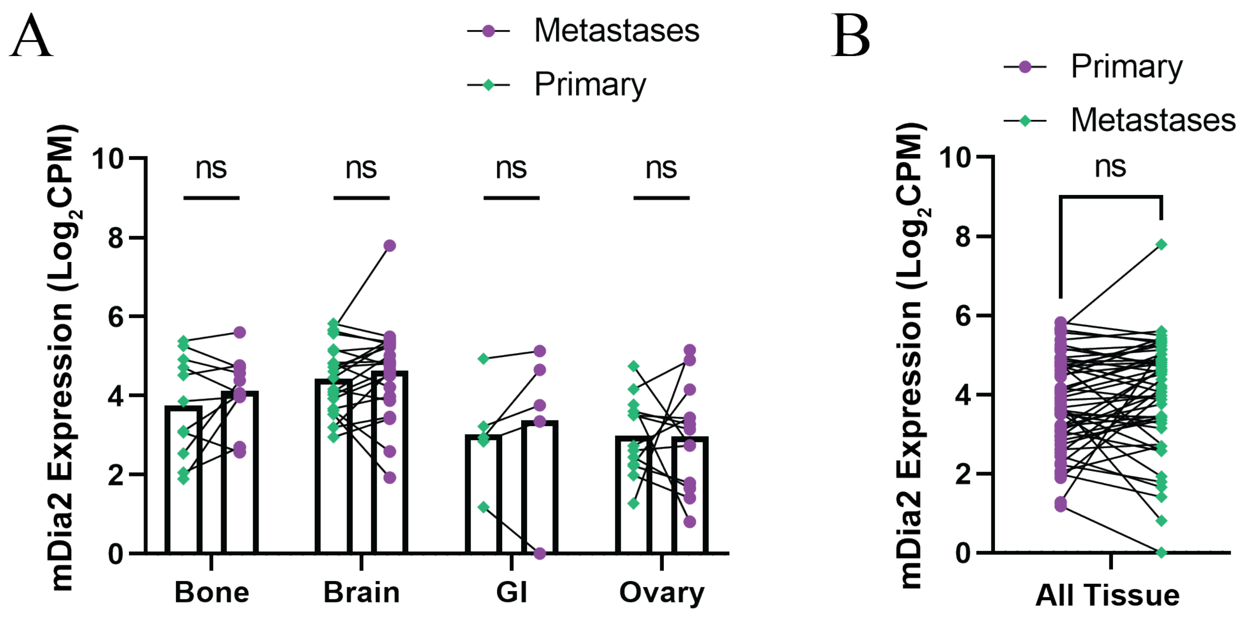

**Figure S8.** Relative mDia2 mRNA expression in pair-matched distant metastasis vs primary tumors in breast cancer patients (*panel A* – organ site-specific display, *panel B* – pooled organ display).

**Table S1.** Table detailing the genes which compose the indicated gene signatures.

| Total MRTF Upregulated Genes | Total MRTF Downregulated Genes | SAP Domain Dependent MRTF Upregulated Genes | SAP Domain Dependent MRTF Downregulated Genes | SRF Domain Dependent MRTF Upregulated Genes | SRF Domain Dependent MRTF Downregulated Genes |
|---|---|---|---|---|---|
| VCAN | KCNAB2 | PPP1R12A | NFKBIA | CCL26 | CD74 |
| TNC | TRIB2 | MYL12A | TBC1D12 | ATOSA | GGT5 |
| SYT1 | CA12 | PLEKHA8 | CRYBG1 | ACTN1 | PLAT |
| MAST4 | PLLP | GARS1 | RAB17 | FERMT2 | GLI3 |
| MCAM | CEMIP | BLVRA | RGS3 | SLC4A4 | SORL1 |
| EDN1 | LFNG | ELL2 | MAL2 | EPB41L4B | CYP1B1 |
| JAG1 | CRTAM | KLHL7 | NIPAL1 | IL11 | HMGA2 |
| TSPAN13 | CTSD | INHBA | ATP9B | MYL9 | MICALL2 |
| RIGI | TNFSF10 | POPDC3 | NNMT | FKBP14 | THBD |
| TPD52L1 | SPINK4 | PTPN22 | CD22 | RASD1 | ALDH1A3 |
| CLIC5 | BMP4 | NIBAN1 | FHL1 | ANXA10 | CSF1 |
| ST8SIA4 | EMP1 | BZW2 | DGAT2 | PTK7 | AHNAK2 |
| QPCT | CDH11 | OSBPL1A | ALDH3A2 | FGF1 | S100A4 |
| DHCR24 | IGFBP4 | HNMT | TXK | RAB32 | S100A2 |
| DNAJC6 | DUSP10 | MAGI1 | PTGS1 | PALLD | TOX |
| KLF7 | CSGALNACT1 | CFAP251 | TGFB1 | DCLK1 | HLA-DRB4 |
| SLC16A7 | ADIRF | HK2 | TIA1 | GSTM3 | APOBEC3G |
| KLHL29 | NR3C2 | MAPKAPK2 | PARD6B | EDNRB | CFB |
| DUSP1 | FBXO32 | RAB8B | CFP | GALNT5 | HLA-DMA |
| SOCS2 | LAPTM5 | PAFAH1B2 | DOCK4 | IGFBPL1 | NFKBIA |
| CALD1 | NPAS2 | ENC1 | CDK15 | TUBB2A | TBC1D12 |
| TEP1 | FAM20C | OXSR1 | ALPP | RAB30 | CRYBG1 |
| AKAP12 | PBX1 | PAWR | FRMD3 | TET1 | RAB17 |
| LCP1 | NANOS1 | NR2F2 | HPSE | GLIPR1 | RGS3 |
| SYTL2 | HLA-DRB3 | LRP10 | MARCKSL1 | GALNT6 | MAL2 |
| PREPL | HLA-DRB1 | LBH | H1-2 | TPM1 | NIPAL1 |
| FGF5 | STMN3 | SPANXB1 | HLA-DMB | CCN1 | ATP9B |
| FRAS1 | HLA-DRB5 | TRAC | | PRKACB | NNMT |
| STON2 | MIR1915HG | LTBP1 | | FLG | |
| TUFT1 | HLA-DPA1 | LAMC2 | | RBMS3 | |
| MYH15 | CEBPD | MXD1 | | COL8A1 | |
| EPHA5 | HLA-DPB1 | P4HA2 | | SLIT2 | |
| TAGLN | HLA-DRA | PHGDH | | GFOD1 | |
| SPOCK1 | CD22 | DSP | | GSN | |
| BANK1 | CD74 | TSPAN1 | | IL18 | |
| LY96 | FHL1 | CDKN1A | | PTPRR | |
| NLRP3 | DGAT2 | IGF2BP3 | | TNIK | |
| EIF4E3 | ALDH3A2 | TLR4 | | RBPMS | |
| PPM1K | TXK | SHC3 | | NEXN | |

| | | | | | |
|---|---|---|---|---|---|
| ZNF23 | PTGS1 | ANKRD1 | | GBP2 | |
| CAVIN2 | GGT5 | GDPD1 | | CPA3 | |
| SMPDL3A | PLAT | LRRC2 | | F2RL2 | |
| KCNQ5 | TGFB1 | SGMS2 | | MATCAP2 | |
| ZNF204P | GLI3 | PGM2 | | PCSK9 | |
| ADGRG1 | TIA1 | TRIB1 | | PLAC1 | |
| VGLL3 | PARD6B | NR2F1 | | PTGER4 | |
| GPX3 | CFP | UPP1 | | BCL2 | |
| KRTAP2-3 | | TACSTD2 | | NEGR1 | |
| **Total MRTF Upregulated Genes** | **Total MRTF Downregulated Genes** | **SAP Domain Dependent MRTF Upregulated Genes** | **SAP Domain Dependent MRTF Downregulated Genes** | **SRF Domain Dependent MRTF Upregulated Genes** | **SRF Domain Dependent MRTF Downregulated Genes** |
| SOGA3 | DOCK4 | HLA-F | | MSRB3 | |
| CCL26 | SORL1 | ZNF542P | | YPEL2 | |
| ATOSA | CYP1B1 | | | RTL3 | |
| LTBP1 | CDK15 | | | ISG15 | |
| LAMC2 | HMGA2 | | | TSC22D2 | |
| MXD1 | ALPP | | | C6orf141 | |
| ACTN1 | MICALL2 | | | FADS3 | |
| P4HA2 | FRMD3 | | | DUSP14 | |
| FERMT2 | HPSE | | | PPP1R12A | |
| SLC4A4 | MARCKSL1 | | | MYL12A | |
| PHGDH | THBD | | | PLEKHA8 | |
| EPB41L4B | ALDH1A3 | | | GARS1 | |
| IL11 | CSF1 | | | BLVRA | |
| DSP | AHNAK2 | | | ELL2 | |
| MYL9 | H1-2 | | | KLHL7 | |
| FKBP14 | S100A4 | | | INHBA | |
| RASD1 | S100A2 | | | POPDC3 | |
| ANXA10 | TOX | | | PTPN22 | |
| PTK7 | HLA-DMB | | | NIBAN1 | |
| FGF1 | HLA-DRB4 | | | BZW2 | |
| TSPAN1 | APOBEC3G | | | OSBPL1A | |
| RAB32 | CFB | | | HNMT | |
| CDKN1A | HLA-DMA | | | MAGI1 | |
| PALLD | NFKBIA | | | CFAP251 | |
| DCLK1 | TBC1D12 | | | HK2 | |
| GSTM3 | CRYBG1 | | | MAPKAPK2 | |
| EDNRB | RAB17 | | | RAB8B | |
| IGF2BP3 | RGS3 | | | PAFAH1B2 | |
| GALNT5 | MAL2 | | | ENC1 | |
| TLR4 | NIPAL1 | | | OXSR1 | |
| IGFBPL1 | ATP9B | | | PAWR | |
| TUBB2A | NNMT | | | NR2F2 | |
| RAB30 | | | | LRP10 | |
| TET1 | | | | LBH | |
| GLIPR1 | | | | SPANXB1 | |
| GALNT6 | | | | TRAC | |
| TPM1 | | | | | |
| CCN1 | | | | | |
| PRKACB | | | | | |
| FLG | | | | | |
| RBMS3 | | | | | |
| COL8A1 | | | | | |
| SLIT2 | | | | | |

| |
|---|
| GFOD1 |
| SHC3 |
| GSN |
| ANKRD1 |
| IL18 |
| PTPRR |
| GDPD1 |
| TNIK |
| RBPMS |

| Total MRTF Upregulated Genes | Total MRTF Downregulated Genes | SAP Domain Dependent MRTF Upregulated Genes | SAP Domain Dependent MRTF Downregulated Genes | SRF Domain Dependent MRTF Upregulated Genes | SRF Domain Dependent MRTF Downregulated Genes |
|---|---|---|---|---|---|
| NEXN | | | | | |
| GBP2 | | | | | |
| CPA3 | | | | | |
| LRRC2 | | | | | |
| SGMS2 | | | | | |
| F2RL2 | | | | | |
| MATCAP2 | | | | | |
| PCSK9 | | | | | |
| PGM2 | | | | | |
| PLAC1 | | | | | |
| PTGER4 | | | | | |
| BCL2 | | | | | |
| NEGR1 | | | | | |
| TRIB1 | | | | | |
| MSRB3 | | | | | |
| YPEL2 | | | | | |
| NR2F1 | | | | | |
| RTL3 | | | | | |
| UPP1 | | | | | |
| TACSTD2 | | | | | |
| ISG15 | | | | | |
| TSC22D2 | | | | | |
| C6orf141 | | | | | |
| HLA-F | | | | | |
| FADS3 | | | | | |
| ZNF542P | | | | | |
| DUSP14 | | | | | |
| PPP1R12A | | | | | |
| MYL12A | | | | | |
| PLEKHA8 | | | | | |
| GARS1 | | | | | |
| BLVRA | | | | | |
| ELL2 | | | | | |
| KLHL7 | | | | | |
| INHBA | | | | | |
| POPDC3 | | | | | |
| PTPN22 | | | | | |
| NIBAN1 | | | | | |
| BZW2 | | | | | |
| OSBPL1A | | | | | |
| HNMT | | | | | |
| MAGI1 | | | | | |
| CFAP251 | | | | | |

| |
|---|
| HK2 |
| MAPKAPK2 |
| RAB8B |
| PAFAH1B2 |
| ENC1 |
| OXSR1 |
| PAWR |
| NR2F2 |
| LRP10 |
| LBH |
| SPANXB1 |
| TRAC |
